# Signal Peptide Peptidase-Like 2b affects APP cleavage and exhibits a biphasic Aβ-mediated expression in Alzheimer’s disease

**DOI:** 10.1101/2022.10.24.513473

**Authors:** Riccardo Maccioni, Caterina Travisan, Stefania Zerial, Annika Wagener, Yuniesky Andrade-Talavera, Federico Picciau, Caterina Grassi, Gefei Chen, Laetitia Lemoine, André Fisahn, Richeng Jiang, Regina Fluhrer, Torben Mentrup, Bernd Schröder, Per Nilsson, Simone Tambaro

## Abstract

Alzheimer’s disease (AD) is a multifactorial disorder driven by abnormal amyloid β-peptide (Aβ) levels. To identify new druggable pathways involved in the Aβ cascade we here investigated the AD pathophysiological role of the presenilin-like intramembrane protease signal peptide peptidase-like 2b (SPPL2b). Aβ42 induced a biphasic modulation of SPPL2b expression in human cell lines and ex vivo mouse brain slices. In addition, SPPL2b was elevated in *App*^*NL-G-F*^ knock-in AD mice as well as in human AD samples. Early high neuronal expression of SPPL2b was followed by a downregulation in late AD pathology in both *App*^*NL-G-F*^ mice and Braak stage V AD brains. Importantly, SPPL2b overexpression or its genetic deletion significantly increased or reduced APP cleavage and Aβ production, respectively. Thus, our results strongly support the involvement of SPPL2b in AD pathology. The early Aβ-induced SPPL2b upregulation may enhance Aβ production in a vicious cycle further aggravating the Aβ pathology suggesting SPPL2b as a potential anti-Aβ drug target.

## INTRODUCTION

Alzheimer’s disease (AD) is a detrimental neurodegenerative condition associated with an abnormal increase of amyloid β-peptide (Aβ) that accumulates in the extracellular space of the brain parenchyma (1) and with an intracellular deposition of neurofibrillary tangles, consisting of hyperphosphorylated tau proteins (2). The levels of Aβ in the brain rise several years before the onset of the disease, promoting the formation of toxic β-sheet oligomers and fibrils that contribute to AD synaptic dysfunction, brain inflammation, and subsequently neurodegeneration (3). Aβ derives from the sequential processing of the amyloid precursor protein (APP) by β- and γ-secretases (4). γ-secretase cleaves APP in the transmembrane region, resulting in the production of Aβ peptides of varying lengths, among which Aβ42 depicts strong neurotoxicity and is highly aggregation-prone (5). Targeting Aβ production and Aβ oligomerization is hence considered a therapeutic approach to counteract or even stop the course of AD (6). Recently, Aβ-immunotherapy showed a positive effect on reducing plaque deposition (7,8) with the monoclonal antibody Aducanumab, which has been approved in the USA by FDA for use in AD patients whereas anti protofibril Aβ antibodies (Lecanemab) reduced cognitive decline (9,10). Still, it is unclear whether this therapeutic approach can block the progression of AD and reverse memory impairment (11).

The use of β- and γ-secretase inhibitors in AD therapy have been unsuccessful, even though, preclinical studies with the same inhibitors showed a substantial reduction in the production of Aβ peptides and plaque deposition in a dose-dependent manner (12,13). In fact, in clinical trials, no positive effects were registered, and, in some cases, the memory impairment was even exacerbated, as well as many side effects were reported (14). β- and γ-secretases are involved in processing over 100 other substrates, in addition to APP, and they are associated with critical physiological pathways such as Notch signaling (14–17). These results highlight the importance of finding more specific and compelling molecules able to lower Aβ and improve memory deficits. For this purpose, a better understanding and characterization of new proteins involved in the Aβ cascade may provide novel, effective AD therapeutic targets.

Signal peptide peptidase (SPP) and SPP-like proteases (SPPLs) are intramembrane proteins that are part of the GxGD type aspartyl protease family including the γ-secretase catalytic part presenilin 1 (PS1) and presenilin 2 (PS2) (18–20). The SPP and SPPL protein family in mammals consists of 5 members: SPP, SPPL2a, SPPL2b, SPPL2c, and SPPL3. In vitro and in vivo characterizations have shown that SPPL2a and SPPL2b share similar substrates such as CD74, LOX-1, and Dectin-1 (21–23). Most importantly, cell-based experiments showed that SPPL2a and SPPL2b cleave the AD-related protein BRI2, TNFα, and the frontotemporal dementia-related protein TMEM106 (24–27). Furthermore, SPPL2b also cleaves the Transferrin receptor-1 which is involved in cellular iron uptake (28). On protein level SPPL2a and SPPL2b have a 40% identity but differ in cellular localization and tissue expression (21,29). SPPL2a is mainly localized in endosomes and lysosomes (30), showing a more ubiquitous expression in all tissues, especially, in immune cells and peripheral tissues (21). In contrast, SPPL2b is localized in the cell membrane and is mainly expressed in the brain, more specifically, in the hippocampus, cortex, and the Purkinje cell layer of the cerebellum (21). The cleavage of the TNFα amino-terminal fragment (NTF) by SPPL2b releases an intracellular fragment, TNFα ICD (25,26) in the cytosol, that promotes the expression of the proinflammatory cytokine interleukin-12 (IL-12). Interestingly, the inhibition of the IL-12 pathway is associated with a reduction of AD pathology and cognitive decline (31). Furthermore, the AD-related SPPL2b substrate, BRI2, is considered an anti-Alzheimer gene (32,33). BRI2 is a 266 amino-acid long transmembrane type II protein strongly expressed in brain neurons and glia (34). BRI2 is cleaved intracellularly by a furin-like protease in its C-terminal region, which results in the release of a 23 amino acid peptide (BRI2_23_) (35). The remaining membrane-bound N-terminal part of BRI2 (mBRI2), which includes the BRICHOS domain, translocates to the cell membrane (36). mBRI2 is further processed by A disintegrin and metalloproteinase domain-containing protein 10 (ADAM10) and subsequently by SPPL2b. While ADAM10 cleaves mBRI2 in its first portion of the extracellular domain, SPPL2b cleaves it in its transmembrane sequence (24). As a consequence, ADAM10 promotes the release of the BRICHOS domain, whereas SPPL2b intermembrane cleavage releases the remaining ectodomain (BRI2-C-peptide) and the intracellular fraction BRI2-ICD (37). In physiological conditions, BRI2 has been reported to negatively regulate Aβ production by binding to APP and inhibiting its processing by the secretases (38,39). We have previously shown in hippocampal primary mouse neurons that BRI2 co-localizes with APP in the soma and dendrites (40). The BRI2 region involved in the interaction with APP consists of the 46-106 amino acids sequence that includes the transmembrane region and the first portion of the extracellular domain (32,41). The APP-BRI2 interaction is interrupted by the SPPL2b-cleavage of BRI2 in the transmembrane region (42). The protective role of BRI2 against Aβ production is also supported by previous findings showing that overexpression of BRI2 reduced the secretion of sAPPα and Aβ peptides *in vitro* and amyloid plaque deposition in an AD mouse model (38,43). Taking this together with the drastically increased level of SPPL2b in the early stage of AD strongly suggests an involvement of this protease in AD pathology (42). Furthermore, this high expression of SPPL2b was correlated with the localization of BRI2-BRICHOS ectodomain in the Aβ plaque deposition and a reduced presence of the APP-BRI2 complexes (42).

Based on these findings, the present study aimed to explore the role of SPPL2b in AD pathogenesis in detail using both cell-based and mouse models, as well as human AD tissues. Accordingly, the pathophysiological role of SPPL2b in Aβ metabolism was evaluated *in vitro* by using human cell lines stably expressing APP, and primary cell cultures from wild-type (WT) and SPPL2b knock-out (KO) mice. Brain slices from WT mice, App knock-in AD mouse model *App*^*NL-G-F*^ and human postmortem AD brain tissues were also used. The *App* gene in *App*^*NL-GF*^ mice contains a humanized Aβ region, APP gene also includes the Swedish “NL”, the Iberian “F,” and the Arctic “G” mutations. *App*^*NL-G-F*^ mice accumulate Aβ and recapitulate several AD-associated pathologies, including Aβ plaques, synaptic loss, microgliosis, and astrocytes in the vicinity of plaques (44). The results obtained support the involvement of SPPL2b in AD pathology and APP processing and suggest that Aβ42 affects SPPL2b expression.

## METHODS

### Human cell cultures

Two human cell lines were used in this study: the human neuroblastoma SH-SY5Y cells both wild-type (SH-SY5Y WT) and stably expressing human APP with the Swedish mutation (SH-SY5Y APPswe); and the human kidney embryonic HEK293 cells WT (HEK293 WT). The cells were cultured with Dulbecco’s modified Eagle’s medium (Gibco™ DMEM) high glucose, supplemented with 10% fetal bovine serum (FBS) and 1% of Penicillin Streptomycin (Gibco) in Petri dishes at 37°C, 5% CO_2_. For the Aβ42 and lipopolysaccharides (LPS) treatments and Western blot analysis, the cells were seeded in 6 well plates at the initial concentration of 300 000 per well and treated after they reach 70 % of confluency. For immune cytochemistry analysis, both cell lines were plated on sterilized glass coverslips positioned into 24-well plates at the initial concentration of 50 000 cells per well. For the HEK293 cells, the coverslips were previously coated with Poly-D-lysine (Sigma). The coverslips were washed three times with PBS 1X at room temperature and fixed in 1 mL/well of PFA 4% (Sigma) under the chemical hood for 10 minutes. Subsequently, coverslips were washed 3 times in PBS 1X and stored in PBS 1X for up to 2 weeks at 4 °C.

### SPPL2b HEK293 transient transfection

HEK293 WT cells were transiently transfected with human SPPL2b plasmid, which has been described earlier (24), using lipofectamine™ 3000 (Invitrogen, L3000008). Cells were plated in a 6-well plate and transfected upon reaching 70-90% confluency. 7.5 μl of lipofectamine™ 3000 Reagent was diluted into 125 μl of Opti-MEM™ Medium and 2.5 μg of DNA vector was diluted into 125 μl of Opti-MEM™ Medium and 5 μl of P3000™ Reagent. The DNA solution was mixed with the Lipofectamine™ 3000 and added to the cells. After 3 days, the cells were collected, and the proteins were extracted for WB analysis.

### SPPL2b Knock-Out cells

The SPPL2b gene was knocked out in HEK293 WT cells by using the SPPL2b CRISPR/Cas9 Knockout Plasmid kit from Santa Cruz Biotechnology® (sc-405646). Briefly, 24 hours before transfection 1.5 × 10^5^ cells were seeded in a 6-wells plate with 3 mL of antibiotic-free medium (DMEM (Gibco™) high glucose) + 10% FBS. On the day of the transfection two separate solutions were prepared: Solution A was made by adding 2 μg of plasmid DNA (Santa Cruz Biotechnology ®, sc-405646) into 130 μl of plasmid transfection medium (Santa Cruz Biotechnology ®, sc-108062); Solution B was made by adding 10 μl of Ultracruz® transfection reagent (Santa Cruz Biotechnology ®, sc-395739) into 140 μl of plasmid transfection medium (Santa Cruz Biotechnology ®, sc-108062). Both solutions were left at RT for at least 5 minutes and were subsequently combined, mixed, and incubated at RT for 20 minutes. The cell media was replaced and the plasmid DNA/Ultracruz® Transfection Reagent Complex (solution A + solution B) was added. Cells were incubated at 37°C, 5% CO_2_ for 72 hours, and the medium was changed at 24 hours post-transfection. Transfected cells were GFP-positive and were selected by fluorescence-activated cell sorting (FACS). The selected cells were plated in a low density in Petri dishes to form cell single colonies. Upon reaching a colony size of approximately 100 cells the single-cell clones were collected using Scienceware Cloning Discs following the manufacturer’s instructions. The cloning discs were soaked in 0.05% trypsin-EDTA for 5 minutes and then placed on top of the marked single-cell colonies. After incubating the cells for 5 min at 37 °C the cells sticking to the cloning discs were transferred into a 24-well plate. When the cells reached confluency, SPPL2b KO cells were validated by ***W***estern blot.

### Mice

*App*^*NL-G-F*^ mice (App^tm3.1Tcs^/App^tm3.1Tcs^) were bred in the animal facility, Solna campus, Karolinska Institutet. SPPL2b KO (B6; CB-*3110056O03Rik*^*Gt(pU-21T)160Imeg*^) mouse embryos were obtained from the Center for Animal Resources and Development at Kumamoto University and rederived in the Karolinska Center for Transgene Technologies (KCTT) Comparative Medicine, at Karolinska Institutet. Mice with the same background (C57BL/6J) were used as controls. Mice were caged in groups of three to five individuals, and the light-dark condition was 12-h:12-h (lights on at 7:00). The ethical permission for described animal experiments has been obtained from Stockholm ethical board (15758-2019). Housing in animal facilities was performed under the control of veterinarians with the assistance of trained technical personnel. All necessary steps to minimize the number of experimental animals were considered. WT and *App*^*NL-G-F*^ female mice 3, 10, and 22 months old were anesthetized with 2% isoflurane and intracardially perfused with PBS. The brains were quickly removed and dissected in two parts. One hemisphere was put in 10 % formalin and later used for morphological and immunohistochemical studies, the other half was dissected, and the hippocampus and cortex were isolated for biochemical analysis. SPPL2a/b double KO mouse brain tissues were obtained from Professor Bernd Schröder (Technical University Dresden, Germany).

### Mouse primary cell culture

Primary cell cultures derived from WT and SPPL2b KO mice were prepared from E17 days embryos. The embryos’ brain tissues were dissected in ice-cold Hank’s balanced salt solution (HBSS, ThermoFisher, #14025092). Dissected tissue was incubated with HBSS supplemented with Accutase (1 ml/brain), centrifuged, and resuspended with Neurobasal medium (Gibco, # 21103049) supplemented with B-27 2 % (Gibco, #17504044) and Glutamax 1 % (Gibco, # 35050061), filtered, and plated in Poly-D-lysine coated wells (Sigma, P6407). At 14 days, the neurons were either collected for Western blot analysis or fixed with formaldehyde 4% for immunofluorescence staining.

For microglia and astrocyte cell cultures, the dissected brain tissues were dissolved in DMEM F12 medium (Gibco, # 21331020) supplemented with 10 % FBS and N2. The cells were filtered and plated in a Petri dish. After 15 days, the astrocytes were separated from microglia cells by mild trypsinization. Briefly, the media was removed, cells were washed in ice-cold PBS, and 5 ml of 0.08% trypsin containing 0.35 mM EDTA (25200-072, Life Technologies, Somerset, NJ) in Dulbecco’s modified Eagle medium (DMEM; 31330-038, Life Technologies) were added to the mixed cultures and incubated at 37 °C. Every 10 min, the Petri media was gently mixed until the astrocyte layer was detaching. The media containing the astrocyte layer was removed and diluted 1:1 with DMEM F12 and centrifuged for 5 minutes at 900 rpm. The pellet was resuspended in DMEM F12 10%FBS, and the cells were plated in 6 or 24 well plates. The remaining pure microglial population was incubated with 0.25% trypsin + 1 mM EDTA for 10 min, dissociated by vigorous pipetting, and resuspended in culture media. After 5 min centrifugation at 2.5 rpm, the cells were resuspended in DMEM/F12 supplemented with 10% FBS and 1% penicillin-streptomycin and plated in 6 or 24 well plates.

### Aβ preparation and treatment

Met-Aβ residues 1–42 (referred to as Aβ42) were recombinantly prepared as described (45). In brief, the Aβ42 peptide was recombinantly expressed in BL21*(DE3) pLysS *Escherichia coli* and purified with DEAE-Sepharose (GE Healthcare). To get rid of large Aβ42 aggregates, the eluted fraction from DEAE-Sepharose (GE Healthcare) was filtered by a 30 000 Da Vivaspin concentrator (GE healthcare) at 4°C and 4000×g. Aβ42 peptides in the filtrate were further concentrated to ∼50 μM at 4°C and 4000×g with a 5 000 Da Vivaspin concentrator (GE Healthcare). The Aβ42 peptide concentration was calculated using an extinction coefficient of 1400 M-1cm-1. Aβ42 peptides were aliquoted in low-bind Eppendorf tubes (Axygene) and stored at - 80 °C.

### Ex Vivo Brain Slices

To study the effect of Aβ42 and the selective glial activator LPS *ex vivo*, acute horizontal WT mice brain slices were obtained as previously described (Andrade-Talavera et al., 2020). Animals were deeply anesthetized with isoflurane (3 per group) and the brain was quickly dissected out and placed in ice-cold artificial cerebrospinal fluid (aCSF) modified for dissection. Slicing aCSF contained (in mM) 80 NaCl, 24 NaHCO_3_, 25 glucose, 1.25 NaH_2_PO_4_, 1 ascorbic acid, 3 Na pyruvate, 2.5 KCl, 4 MgCl_2_, 0.5 CaCl_2_ and 75 sucrose and bubbled with carbogen (95% O_2_ and 5% CO_2_). Horizontal sections (350 μm thick) of both hemispheres were prepared with a Leica VT1200S vibratome (Leica Microsystems, Wetzlar, Germany). Immediately after cutting, slices were transferred into a humidified interface holding chamber containing standard aCSF (in mM): 124 NaCl, 30 NaHCO_3_, 10 glucose, 1.25 NaH_2_PO_4_, 3.5 KCl, 1.5 MgCl_2_, and 1.5 CaCl_2_. The recovery chamber was continuously supplied with humidified carbogen, and slices were allowed to recover for a minimum of 1 h before any incubation was performed. Next, slices were incubated for 6 hrs. with 50 nM Aβ42, 20ug/ml LPS, or standard aCSF continuously bubbled with carbogen. After incubation cortical and hippocampal sections were snap-frozen for Western blot analysis determinations.

### Human tissues

Post-mortem brain tissues were obtained from Netherlands Brain Bank (Amsterdam, Netherlands), and SPPL2b immunoblotting was performed in brain homogenates from the Netherlands Brain Bank under ethical permit 935S and from Karolinska Institutet. AD cases had a clinical diagnosis of sporadic AD during life and fulfilled post-mortem neuropathological consensus criteria for AD.

### Western Blot

Cells and brain tissues were submerged in RIPA buffer (ThermoFisher) containing phosphatase (phosphatase inhibitors cocktails, Sigma) and protease inhibitors (mammalian protease arrest, G biosciences). Cell and tissue suspensions were homogenized, sonicated, and centrifuged at 14,000 g for 20 minutes at 4°C. Supernatants were collected and the protein concentration was calculated. The samples were stored at -80°C until use. The protein concentration was measured using the Pierce™ BCA Protein Assay kit. Samples, 20 or 50 μg of protein per well, were separated by Mini-PROTEAN® TGX™ Precast Gels 4-20% (Bio-Rad) and transferred to nitrocellulose membranes (Bio-Rad). The transfer was performed with Blot Turbo (Bio-Rad) for 30 min at 25V. Membranes were blocked in 5% milk in Tris-Buffered Saline 0.05% Tween 20 (TBS-T) for 1 h at room temperature or overnight at 4°C.

The membranes were incubated with the primary antibodies diluted in TBS-T for 2 h at room temperature or overnight at °4C (see antibodies dilutions in Supl. Table1). After washing the membranes were incubated in fluorescently labeled secondary antibodies (LiCor) for 1 hour at room temperature.

### ELISA

Secreted Aβ40 and Aβ42 peptides in cell culture media were measured using the human amyloid-β (1-40) or the human amyloid-β (1-42) ELISA kits from Immuno-Biological Laboratories (IBL-27711; IBL-27713). The absorbance was measured at 450 nm by a Microplate reader (Multiskan SkyHigh, ThermoFisher). The protein concentration was calculated by using a standard curve as reported by the manufacturer.

### Immunofluorescence (IF)

The cells were fixed with 4% PFA and washed 3 times in DPBS 1X after they were permeabilized using 0.1% Triton-X100/PBS 1X for 10 minutes and then washed 3 times with PBS 1X. The blocking was performed by incubating the cells with 3% BSA/PBS 1X for 1 hour at room temperature. After the 3 additional washing steps were performed and incubated overnight at 4 °C with the primary antibody in 3% BSA/PBS 1X. The day after the cells were washed and incubated with the secondary antibody diluted 1:1000 for 1 hour at room temperature. After the washing, the cells were incubated in Hoechst solution (Hoechst 3342 – Thermo Scientific ™) with a dilution of 1:500 for 15 minutes at room temperature. A final step of washing was performed and then glasses were mounted using Fluoroshield™ histology mounting medium (Sigma-Aldrich.) on Superfrost™Plus Adhesion Microscope Slides (Epredia). The fluorescence intensity between cells was calculated by using ImageJ software (National Institutes of Health, MD). The total cell fluorescence (TCF) was calculated using this formula; TCF= Integrated Density – (Area of selected cell X Mean fluorescence of background readings.

Paraffin-embedded brain tissues were sectioned into 5 μm thick sections. The tissue sections were put on glass slides and deparaffinized by washing in Xylene and decreasing concentrations of ethanol (99-70%). For antigen retrieval, slides were pressure boiled in citrate buffer solution (0.1 M citric acid and 0.1 M sodium citrate) at 110°C for 5 min and then washed with tap water followed by PBS-Tween 0.05% for 5 minutes each. Sections were then incubated with TNB blocking buffer (0.1M Tris-HCl pH 7.5, 0.15M NaCl, and 0.5% Blocking Reagent; PerkinElmer, USA) or NGS (normal goat serum, Vector Laboratories, USA) for 30 min at room temperature. The brain sections were then incubated with the primary antibody at 4°C, overnight (see antibodies dilutions in Supl. Table1). Thereafter, sections were incubated with biotinylated anti-mouse or anti-rabbit antibodies (Vector Laboratories; UK) 1:200 in TNB buffer or NGS for 2 hours at room temperature and then incubated with HRP-Conjugated Streptavidin (PerkinElmer; USA) 1:100 in TNB buffer or NGS for 30 min. For signal amplification, samples were incubated for 10 min in tyramide (TSA PerkinElmer; USA) 1:50 in Amplification Reagent. Finally, samples were incubated for 15 min with slow agitation with Hoechst solution, 1:3000 in PBS-T followed by mounting with PermaFluor Aqueous Mounting Medium (ThermoScientific, USA) and kept for drying overnight. Between each incubation step, samples were washed 3x in PBS-T for 5 min with slow agitation. The sections were then visualized with Nikon Eclipse E800 confocal microscope and imaged with a Nikon DS-Qi2 camera in 2x, 10x and 20x magnifications. Quantification of the immunoreactive signals was performed using ImageJ software (National Institutes of Health, MD); in the cortex region the total number of positive cells stained was counted, in the hippocampus, the fluorescence intensity was calculated in the region of interest (Cornu Ammonis 3; CA3).

### Immunohistochemical (IHC)

Coronal mouse brain sections were de-paraffinized and re-hydrated as reported previously for IF. Sections were pressure boiled in a Decloaking Chamber (Biocare Medical) immersed in DIVA decloaker 1X solution (Biocare Medical, Concord, USA) at 110 °C for 5 min. Slides were let cool down at RT for 20 min, then washed with PBS buffer containing 0.1% tween 20 (PBST) and incubated with peroxidase blocking solution (Dako) for 5 min. The sections were washed in Tris-buffered saline (TBS) and additional blocking was performed with a Background punisher (Biocare) for 10 min. Primary antibodies diluted in DAKO (Agilent) antibody diluent were incubated for 45 min at RT. Slides were then washed in TBS and incubated with Mach 2 Double stain 2 containing alkaline phosphatase (AP) conjugated secondary anti-rabbit antibody for 30 min at RT. AP staining was detected with permanent red (Biosite). Sections were counterstained with hematoxylin (Mayer), de-hydrated through ethanol (from 70% to 99%), cleared in xylene, and mounted with DEPEX mounting media (Merck).

### Statistical analysis

All statistical comparisons were performed using Prism 8 (GraphPad Software Inc., CA, USA). Results were analyzed by multiple t-tests or unpaired Student’s *t*-test. For comparisons among more than three groups, one-way ANOVA was used followed by Tukey’s multiple comparison test. Variability of the estimates was reported as the standard error of the mean (SEM), and *p*<0.05 was considered statistically significant for all the analyses.

## RESULTS

### SPPL2b is up-regulated in human SH-SY5Y cells overexpressing APP with the Swedish mutation

The expression levels of SPPL2b in healthy and AD-like pathology were initially evaluated in the neuronal-like human neuroblastoma cell line SH-SY5Y and the human kidney cell line HEK293. Both WT cells (SH-SY5Y WT; HEK293 WT), and SH-SY5Y cells stably overexpressing human APP with the Swedish mutation (SH-SY5Y APPswe) were used (Fig.S1A). SH-SY5Y APPswe contains a substitution of two amino acids, lysine (K) and methionine (M) to asparagine (N) and leucine (L) (KM670/671NL), resulting in a 10- to 50- fold higher processing of APP by the β-secretase to and an increased production Aβ40 and Aβ42.

APP overexpression in the SH-SY5Y-APPswe cell induced an increase of SPPL2B revealed by a specific SPPL2b band appearing at 70kDa in Western blot analysis, which was confirmed by using two different antibodies that recognize the N-terminal (93-142aa, extracellular) and the C-terminal (541-592aa, intracellular) regions respectively of the SPPL2b protein (Fig.S1B-E). Further, Crispr/Cas9-generated SPPL2b ko HEK293 WT cells were used as negative controls (Fig.S1D).

Notably, Western blot analysis showed that overexpression of APPswe in SH-SY5Y leads to 4 times increase of SPPL2b compared to WT cells (Fig. 1A, B), which was further confirmed by IF staining (Fig. 1A). The higher SPPL2b expression in SH-SY5Y APPswe correlates with increased secretion of Aβ40 and Aβ42 in the conditioned media as compared with SH-SY5Y WT media (Fig 1C, D).

**Figure 1.**
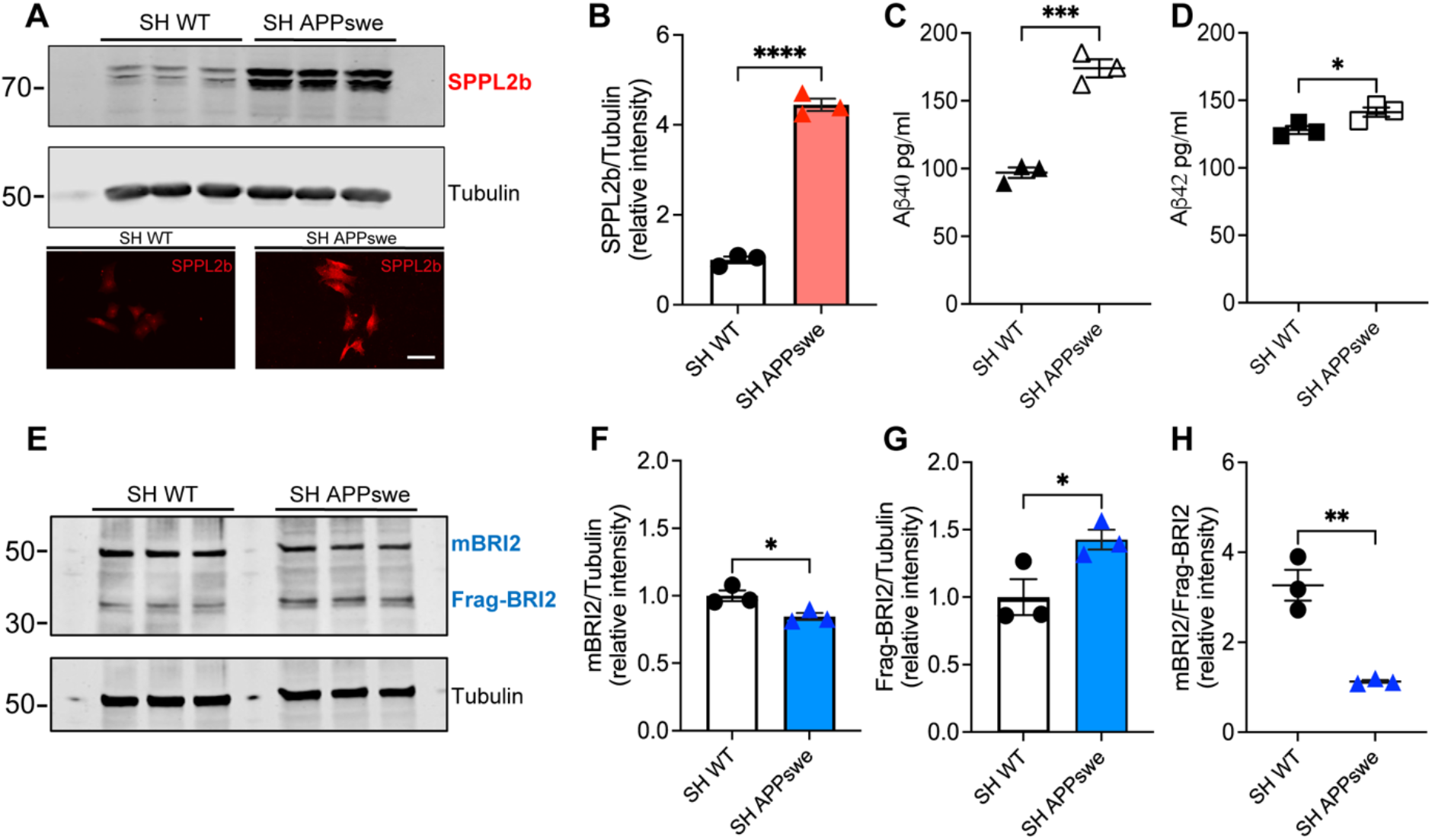
SPPL2b is up-regulated in SH-SY5Y APPswe cells. (**A**) Western blot and immunofluorescence analysis of SPPL2b (Invitrogen, PA5-42683) expression in SH-SY5Y WT (SH WT) and SH-SY5Y APPswe (SH APPswe) cells. (**B**) Quantification of the SPPL2b/Tubulin ratio in the Western blot from A. ELISA analysis of Aβ40 (**C**) and Aβ42 (**D**) concentration in the media of SH-SY5Y WT and SH-SY5Y APPswe cells. (**E**) Western blot analysis of BRI2 (goat Anti-Bri2 BRICHOS antibody) expression in SH-SY5Y WT and APPswe cells. (**F**) Quantification of mature BRI2 protein (mBRI2, 50kDa) expression normalized with tubulin protein expression. (**G**) Quantification of the BRI2 cleavage fraction (Frag-BRI2, 35kDa), (**H**) and quantification of the BRI2 50kDa/35kDa ration in the Western blot in **E**. Data are represented as mean ± S.E.M. *P<0,05; **P<0,01; ***P<0,001 significantly different from SH-SY5Y WT. Data were analyzed by unpaired Student’s t-test.

In addition to the higher SPPL2b expression, Western blot analysis revealed a significant reduction of mature BRI2 (at 50kDa band) in SH-SY5Y APPswe cells, and an increase of BRI2 proteolytic fragment containing the BRICHOS domain (Frag-BRI2; at 35kDa band) (Fig. 1E-G) as compared to SH-SY5Y WT cells. Consequently, the ratio of mBRI2 and Frag-BRI2 was significantly reduced in the SH-SY5Y APPswe cells (Fig. 1H). However, previous *in vitro* findings reported that the cleavage of mBRI2 by SPPL2b happens with a higher efficiency after a pre-cleavage from ADAM10 (24,46); nevertheless, our data support that increased APP/Aβ levels in the SH-SY5Y induce a higher SPPL2b expression and an altered BRI2 secretion and proteolysis.

### Aβ42 affects SPPL2b expression

To analyze whether the increased expression of SPPL2b in the SH-SY5Y APPswe cells is Aβ mediated, SH-SY5Y cells were treated with human recombinant AD-associated Aβ42 (10-100 nM). A biphasic Aβ42 dose-dependent response effect was observed on SPPL2b expression in SH-SY5Y WT cells with an up-regulation at 50 nM and a down-regulation at 100 nM of Aβ42 (Fig. 2A, B). A similar effect was also observed in HEK293 cells (Fig.S2A-D).

**Figure 2.**
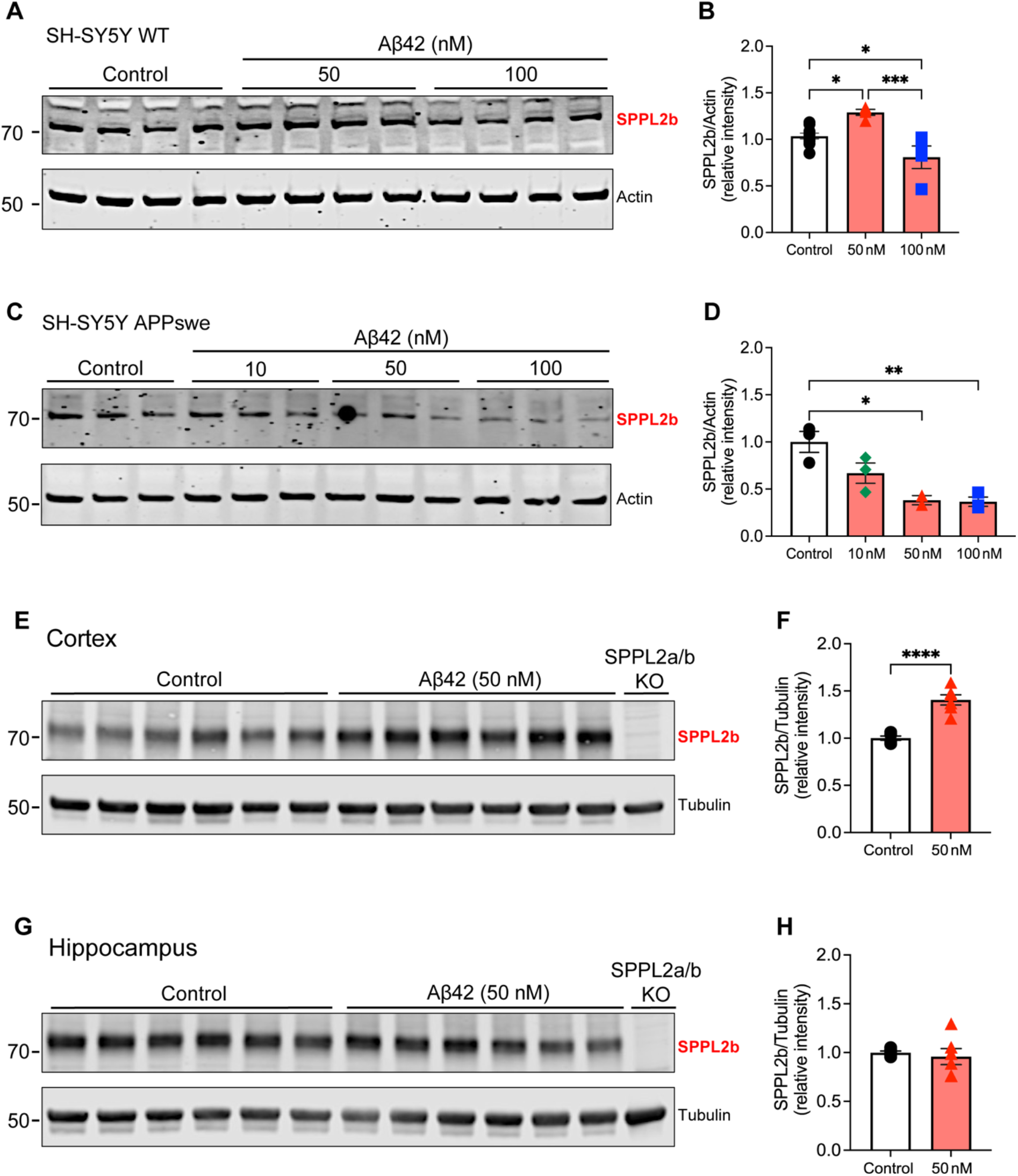
Aβ42 affects SPPL2b expression. Western blot analysis of SPPL2b (Invitrogen, PA5-42683) expression in SH-SY5Y WT (**A, B**) and SH-SY5Y APPswe cells (**C, D**) after 6 hours exposure to increasing concentrations of Aβ42 (10 nM, 50 nM, 100 nM). Results are normalized to actin protein expression. Representative Western blot analysis of SPPL2b (rabbit anti-SPPL2b) expression in mice brain cortex (**E, F**) and hippocampal slices (**G, H**) kept ex vivo in artificial CSF and treated with Aβ42 50 nM for 6 hours. Results are normalized to tubulin protein expression. Data are represented as mean ± S.E.M. ****P < 0.0001, ***P < 0.001, **P < 0.01, *P < 0.05 significantly different from the Control. Data were analyzed either by unpaired Student’s t-test (F, H) or by one-way ANOVA with Tukey’s multiple comparison test (B, D).

On the other hand, when SH-SY5Y APPswe cells were treated with Aβ42, a significant reduction of SPPL2b was observed at the doses of 50 and 100 nM as compared to the untreated cells (Fig. 2. C, D). This phenomenon could be related to a synergic consequence of the presence of both exogenous Aβ42 and the Aβ40/Aβ42 released from SH-SY5Y APPswe cells in the media. Furthermore, we also investigated if SPPL2b expression can be affected by inflammation. To answer this question SH-SY5Y cells were treated with LPS (1 μg/ml), but no changes in SPPL2b expression were observed (Fig.S2 E, F).

Interestingly, similar to the results observed in the SH-SY5Y WT cells, 50 nM of Aβ42 induced a strong up-regulation of SPPL2b in the cortex of acute brain sections from WT mice-maintained ex vivo (Fig. 2E, F). However, this upregulation was not observed in the hippocampus (Fig. 2G, H). As in SH-SY5Y cells, we sought to verify whether inflammation could mediate the expression of SPPL2b in the *ex vivo* preparation. For that purpose, acute brain sections from WT mice were treated with LPS. Interestingly, 20 μg/ml of LPS did not affect the expression of SPPL2b in the cortex (Fig.S2 G, H), but induced a significant increase of SPPL2b in the hippocampus (Fig.S2 I, J), Taken together, these results suggest that Aβ42 might modulate SPPL2b expression in a biphasic manner.

### Overexpression and genetic deletion of SPPL2b affect APP cleavage and Aβ production

After having found that Aβ42 induces an increase in SPPL2b expression we wanted to know whether SPPL2b itself affects the processing of APP and subsequent Aβ generation. To investigate this, we transiently overexpressed human SPPL2b in HEK293 cells (Fig. 3A). Intriguingly, in accordance with what was observed in SH-SY5Y APPswe cells, an altered processing of BRI2 was also observed in HEK293 cells overexpressing SPPL2b (Fig. 3B). Western blot analysis showed an increased level of the soluble BRI2 BRICHOS-containing fragment Frag-BRI2 and a significant reduction of the mBRI2/Frag-BRI2 ratio (Fig. 3B) compared to the WT control cells. In this regard, Western blot analysis of conditional media from the HEK293 overexpressing SPPL2b cells showed the presence of a soluble BRI2 fragment (sFrag-BRI2) by using an anti-BRI2 BRICHOS antibody but not with an anti-BRI2 that recognizes the intracellular domain (Fig. 3C). These data support altered BRI2 processing in the SPPL2b overexpressing cells.

**Figure 3.**
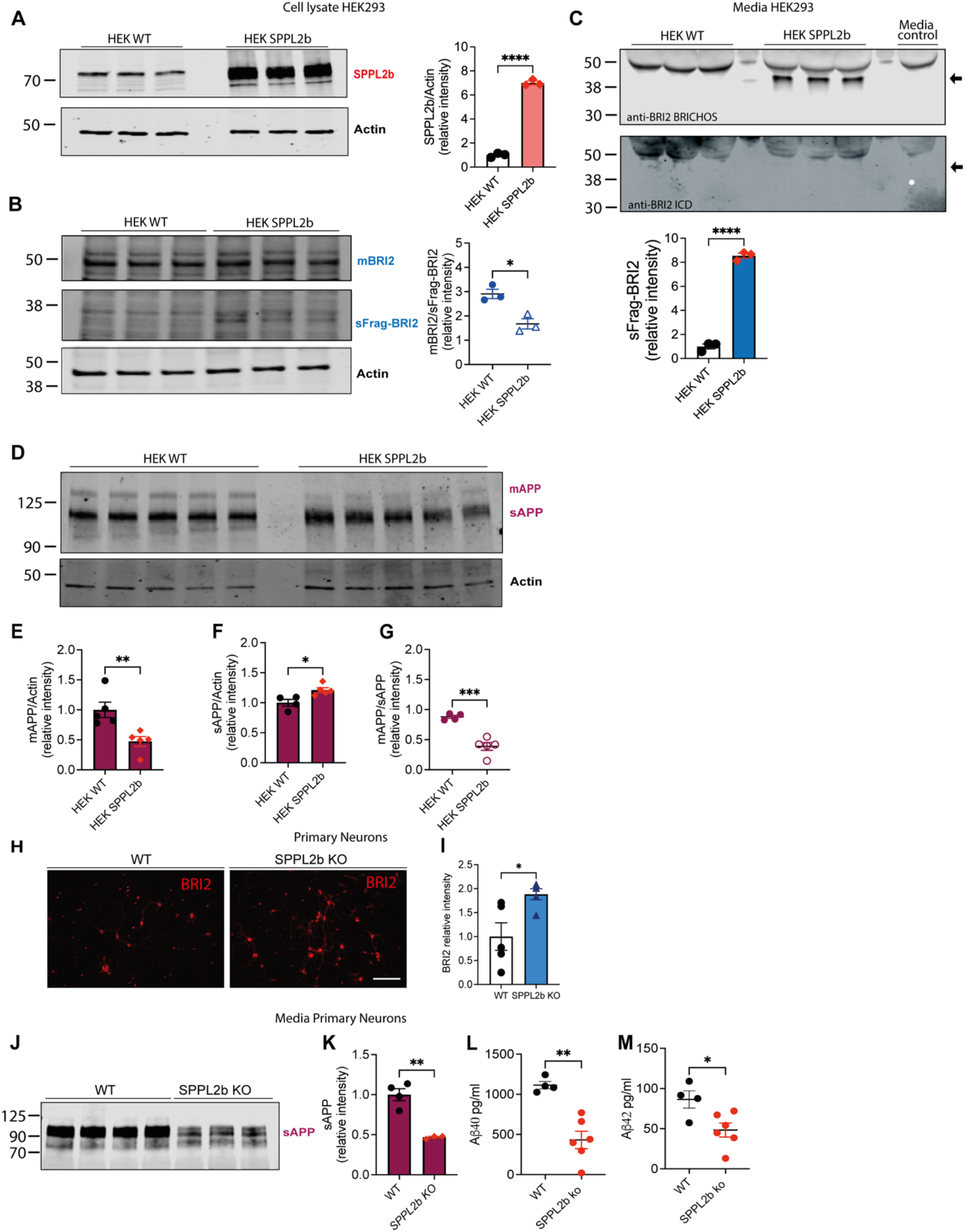
SPPL2b affects APP cleavage. (**A**) Representative Western blot showing SPPL2b expression in HEK293 control cells (HEK WT) and HEK293 cells transiently overexpressing human SPPL2b (HEK SPPL2b) together with the quantification of the Western blot shown. (**B**) Western blot of BRI2 (goat Anti-Bri2 BRICHOS antibody) expression in HEK WT vs HEK SPPL2b cells; quantification of the ratio between BRI2 50 kDa (mBRI2) and 35 kDa (Frag-BRI2) from the Western blot. (**C**) Western blot of BRI2 (goat Anti-Bri2 BRICHOS antibody) in HEK WT vs HEK SPPL2b cell media and its quantification. The arrows indicate the position of the soluble BRI2 band location (sFrag-BRI2). (**D**) Western blot analysis of cellular APP protein levels in lysates from HEK WT and HEK SPPL2b cells; mature (mAPP) and soluble (sAPP) forms of APP were detected with the 22C11 antibody. (**E**, and **F**) Quantitative analysis of the mAPP and sAPP intensity normalized with actin. (**G**) Quantification of the ratio between mAPP and sAPP of the Western blot in (E). (**H**) Immunohistochemistry of BRI2 (Anti-ITM2B Antibody (C-8), Santa Cruz) in cultured mouse primary neurons from WT and SPPL2b KO embryos and (**I**) quantification of BRI2 intensity. Scale bar, 100 μm. (**J**, and **K**) Western blot analysis of sAPP in conditioned media from WT and SPPL2b KO cultured mouse primary neurons and its quantification. (**L**, and **M**) Aβ40 and Aβ42 quantification by ELISA in conditioned media from WT and SPPL2b KO cultured mouse primary neurons. Data are represented as mean ± S.E.M. ****P < 0.0001, ***P < 0.001, **P < 0.01, *P < 0.05 significantly different from the controls. Data were analyzed by unpaired Student’s t-test.

Most importantly, overexpression of SPPL2b also led to a significant reduction of mature APP (mAPP), as compared to the control non-transfected cells (Fig. 3 D, E) concomitant with, a significant increase of soluble APP (sAPP) (Fig. 3 D, F) (47). Consequently, the ratio of mAPP/sAPP was significantly reduced in the SPPL2b overexpressing cells (Fig. 3G). These data reveal that SPPL2b overexpression leads to an increased processing/secretion of both BRI2 and APP.

Moreover, to further verify the modulatory effect of SPPL2b on the APP cleavage process in a neuronal setting we established a primary neuronal cell culture derived from the SPPL2b KO mice. We observed a significant increase in BRI2 staining by using an anti-BRI2-ICD antibody as compared with the control WT neurons (Fig. 3H, I). Most importantly, sAPP was significantly reduced to about 50% in the media of SPPL2b KO neurons (Fig. 3 J, K). Consistently, the quantification of Aβ40 and Aβ42 peptides in the cell media by ELISA showed a significant decrease in both Aβ40 and Aβ42 levels (Fig. 3L, M). Taken together, these data strongly support that SPPL2b affects APP proteolysis and consequently also Aβ generation/secretion.

### SPPL2b follows a biphasic expression upon the progression of Aβ pathology in *App*^*NL-G-F*^ mice

To investigate how Aβ pathology affects SPPL2b levels *in vivo*, SPPL2b expression was evaluated in the *App*^*NL-G-F*^ knock-in AD mouse model. We investigated the expression of SPPL2b at three different AD stages corresponding to early, mid, and late stages of the pathology (respectively 3 months, 10 months, and 22-24 months) (Fig. 4). Interestingly, a higher expression level of SPPL2b was observed in the cortex in the early stage of the AD-associated Aβ pathology in 3 months old *App*^*NL-G-F*^ mice (Fig. 4A), but no significant differences were observed at the same age in the hippocampus area (Fig. 4B). However, at 10 months of age, when Aβ pathology is severe and accompanied by neuroinflammation in this mouse model, SPPL2b protein expression was significantly lowered in both cortex (Fig. 4C) and the hippocampus (Fig. 4D) as compared to age-matched control mice. A significant down-regulation of SPPL2b expression was also observed in the hippocampus and cortex in a very late stage of the pathology (22-24 months of age) in the *App*^*NL-G-F*^ mice as compared to WT mice (Fig. 4E, F). These results support the hypothesis of a biphasic modulation of SPPL2b in *App*^*NL-G-F*^ mice.

**Figure 4.**
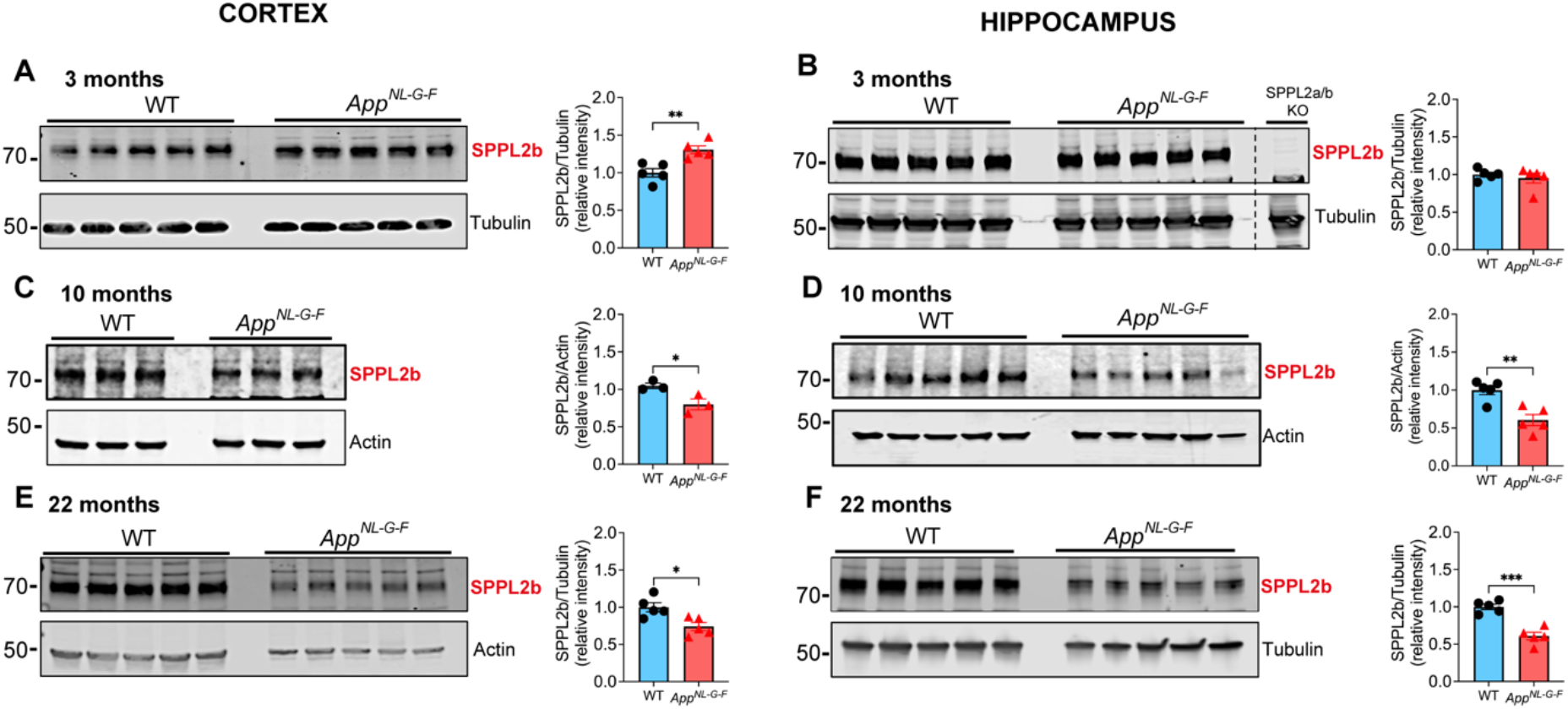
Early high expression of SPPL2b is followed by a downregulation in the late AD-associate Aβ pathology in *App*^*NL-G-F*^ mice. Western blot analysis of SPPL2b expression in WT and *App*^*NL-G-F*^ cortex and hippocampus at 3 months (**A, B**), 10 months (**C, D**), and 22 months (**E, F**) (n= 3 to 5 for each time point). In the Western blot in B, in the last lane, was used a mouse hippocampus SPPL2a/b KO samples as a negative control. SPPL2b protein levels were normalized by using β-actin or tubulin as a loading control. Data are represented as mean ± S.E.M. ****P < 0.0001, ***P < 0.001, **P < 0.01, *P < 0.05 significantly different from the WT mice. Data were analyzed by unpaired Student’s t-test.

### SPPL2b is mainly expressed in neurons and microglia deposited in the amyloid plaques

SPPL2b expression in *App*^*NL-G-F*^ and WT mice was further evaluated by immunofluorescence (Fig. 5). The quantification of SPPL2b-positive cells confirmed the decrease of SPPL2b in the cortex and hippocampus in *App*^*NL-G-F*^ mice at 10 and 22 months of age (Fig. 5A, B) as shown by the Western blot analysis (Fig. 4). In particular, the retrosplenial and the auditory cortex areas exhibited intense staining in the WT mice but were also the most affected in the *App*^*NL-G- F*^ mice with a specific decrease in the layer I-V (Fig.S3C). Notably, the retrosplenial cortex is implicated in navigation and contextual memory, the auditory cortex, instead, is specialized for processing speech sounds and other temporally complex auditory signals (48,49). Interestingly, both areas are impaired in AD (50,51).

**Figure 5.**
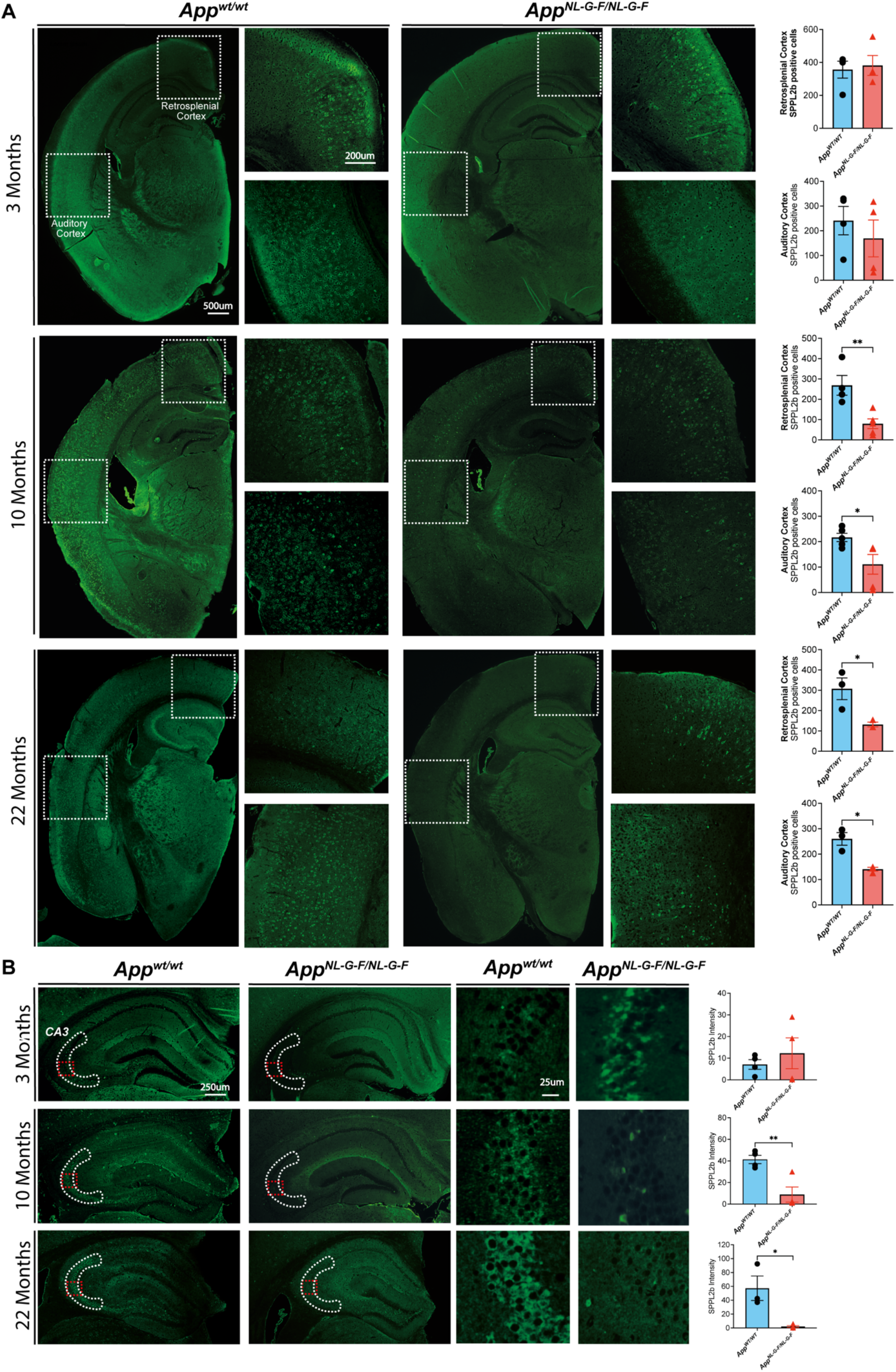
SPPL2b staining in *App*^*NL-G-F*^ mice 3, 10, and 22 months old. (**A**) Representative positive SPPL2b staining in 3-, 10-, and 22months old WT and *App*^*NL-G-F*^ mice. The dashed squares indicate the retrosplenial and the auditory cortex regions. On the quantification of SPPL2b positive cells in the retrosplenial and the auditory cortex regions. (**B**) SPPL2b staining in the hippocampal area in 3, 10, and 22months old *App*^*NL-G-F*^ mice. The white dashed area highlights the CA3 hippocampal area, the region of interest (ROI) analyzed. The red dashed squares indicate the magnified area reported on the right. Data are represented as mean ± S.E.M. of 3 to 5 mice from each group. Data were analyzed by unpaired Student’s t-test. Scale bar sizes are reported in the figure.

Furthermore, the quantification of the hippocampal SPPL2b staining showed also a significantly lowered SPPL2b staining of CA1 pyramidal neurons in the cornu ammonis 3 area (CA3) of *App*^*NL-G-F*^ as compared to the WT mice (Fig 5B). The CA3 region has an important role in memory processes, especially at the initial stage of acquisition (52).

Interestingly, in10-month-old *App*^*NL-G-F*^ mice with an established Aβ pathology, SPPL2b positive staining was additionally found in the proximity of Aβ plaques (Fig. 6A). To identify the cellular origin of this SPPL2b staining we evaluated the expression levels of SPPL2b in neurons and in glial cells in WT and *App*^*NL-G-F*^ mice by performing double staining with neuronal (NeuN), microglia (Iba1) and astrocyte (GFAP) markers. We found that SPPL2b protein is expressed in neurons, especially in layer I-V in the cortex (Fig. 6B), and in glia surrounding the amyloid plaque (Fig. 6C), data that further support the involvement of SPPL2b in AD pathology. No colocalization between SPPL2b and the astroglia marker GFAP was detected (Fig. 6D).

**Figure 6.**
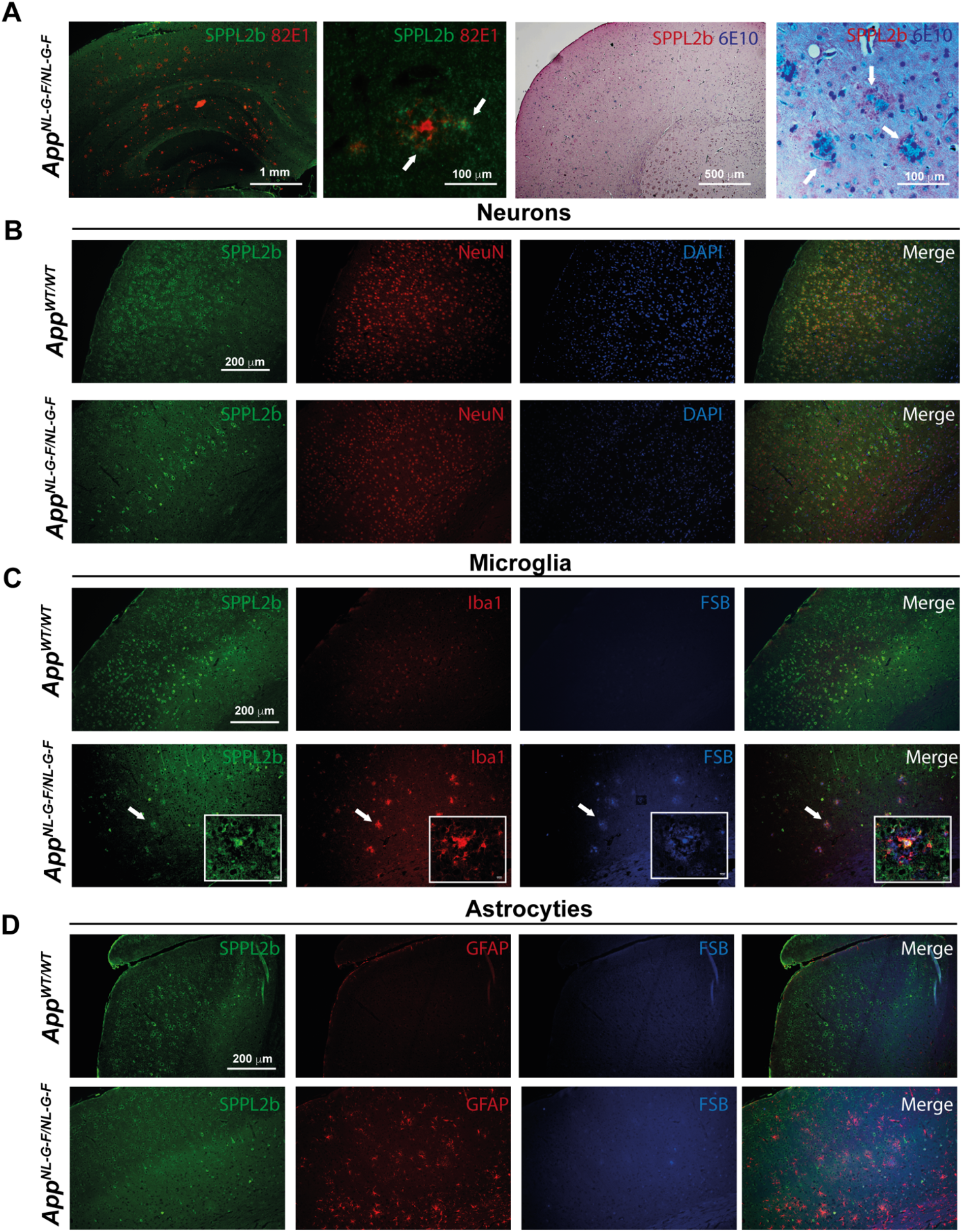
SPPL2b is mainly expressed in neurons and microglia associated with amyloid plaques. **(A)** Representative immunofluorescence and immunohistochemistry staining of SPPL2b surrounding Aβ plaques in *App*^*NL-G-F*^ mice 10 months old by using the anti-amyloid antibodies 82E1 and 6E10 respectively. (**B**) SPPL2b protein staining in neurons, (**C**) in microglial, and (**D**) astrocytes. In green, the staining of SPPL2b; in red, the staining of neurons (NeuN), microglia (Iba1), and astrocyte (GFAP); in blue, the nucleus (DAPI) and Aβ plaques (FSB). Scale bar sizes are reported in the figure.

### Aβ42 modulates SPPL2b expression in neuronal and glial cells in opposite directions

SPPL2b expression was further evaluated in neuronal and glial mouse primary cell culture in both physiological conditions and after exposure to Aβ42. A strong SPPL2b staining was observed in the neuronal soma and in the neurites (Fig. 7). Interestingly, SPPL2b was also found in the dendritic spines (Fig. 7A). Furthermore, a colocalization of SPPL2b with APP was observed in the neuronal soma (Fig. 7A). These findings are in line with the IF results from the mouse brain, in which intense staining in the neuronal soma was also observed. A significant down-regulation of the SPPL2b staining was observed in neuronal cells treated with Aβ42 (1 μM) for 24 hours (Fig. 7B). This reduction in SPPL2b protein levels is in line with the data observed in the 10 and 22 months old *App*^*NL-G-F*^ mice, where a high Aβ pathology is present. This finding further supports the involvement of Aβ42 in modulating the SPPL2b expression. A positive SPPL2b staining was also observed in microglia primary cell culture. However, in contrast to what we observed in neurons, the exposure to Aβ42 for 24 hours increased SPPL2b expression in microglia cells. These phenomena correlate with a higher Iba1 staining and microglia activation (Fig. 7C). Finally, astroglia showed a low level of SPPL2b staining which was not affected by Aβ42 exposure (Fig.S4). Western blot analysis confirmed the presence of SPPL2b in neuronal and glial cells, where astrocytes have a 3.7 times lower expression than neurons (Fig.S4D, E).

**Figure 7.**
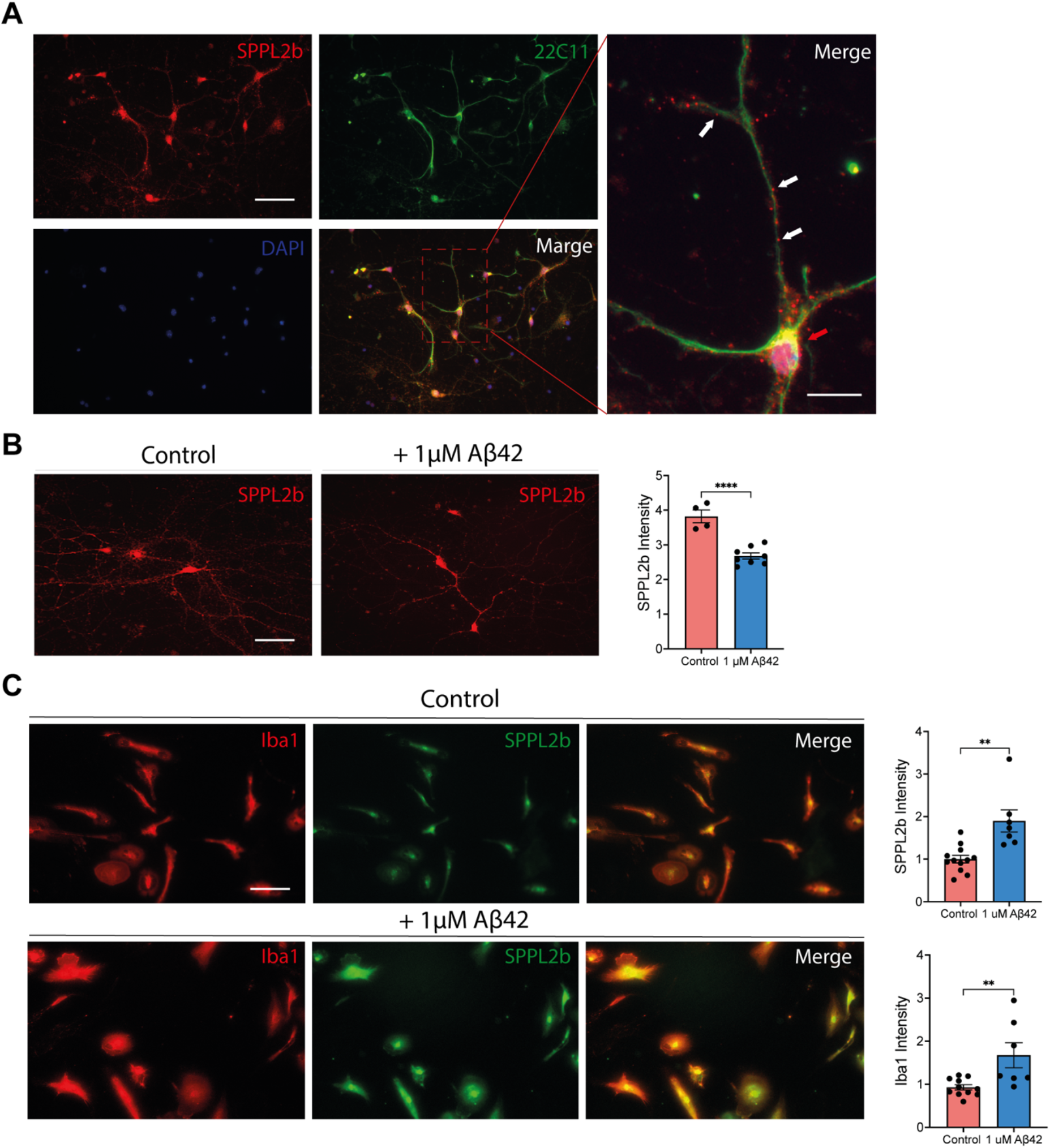
SPPL2b expression in primary neurons and microglia. (**A**) Representative SPPL2b, APP (22C11), nucleus (DAPI), and merge staining in primary neuronal cell culture of WT embryos. On the right, the red arrow highlights SPPL2b-APP colocalization in the neuronal soma area. The white arrow indicates SPPL2b positive staining in neuronal spines. (**B**) SPPL2b expression in control conditions and after a 24-hour treatment with 1μM Aβ42 and quantification (n=4 to 8 slices) of the stainings. Data are represented as mean ± SEM. ****P<0,0001 vs WT. (**C**) SPPL2b expression was monitored in primary microglial cell culture was not treated or treated with 1μM Aβ42 for 24 hours. The microglia marker Iba1 is in red, and SPPL2b in green. Quantifications of SPPL2b and Iba1 staining are shown on the right. Data are represented as mean ± S.E.M. ****P < 0.0001, **P < 0.01 significantly different from the controls. Data were analyzed by unpaired Student’s t-test. Scale bar sizes are reported in the figure.

### SPPL2b is downregulated in AD human prefrontal cortex

Intriguingly, and in agreement with the results obtained in aged *App*^*NL-G-F*^ mice (10 and 22 months of age), a Western blot study of postmortem AD human prefrontal cortex from patients in the late stage of the disease (Braak stages V-VI) exhibited a significant SPPL2b decrease as compared to non-AD samples (Fig. 8A).

**Figure 8.**
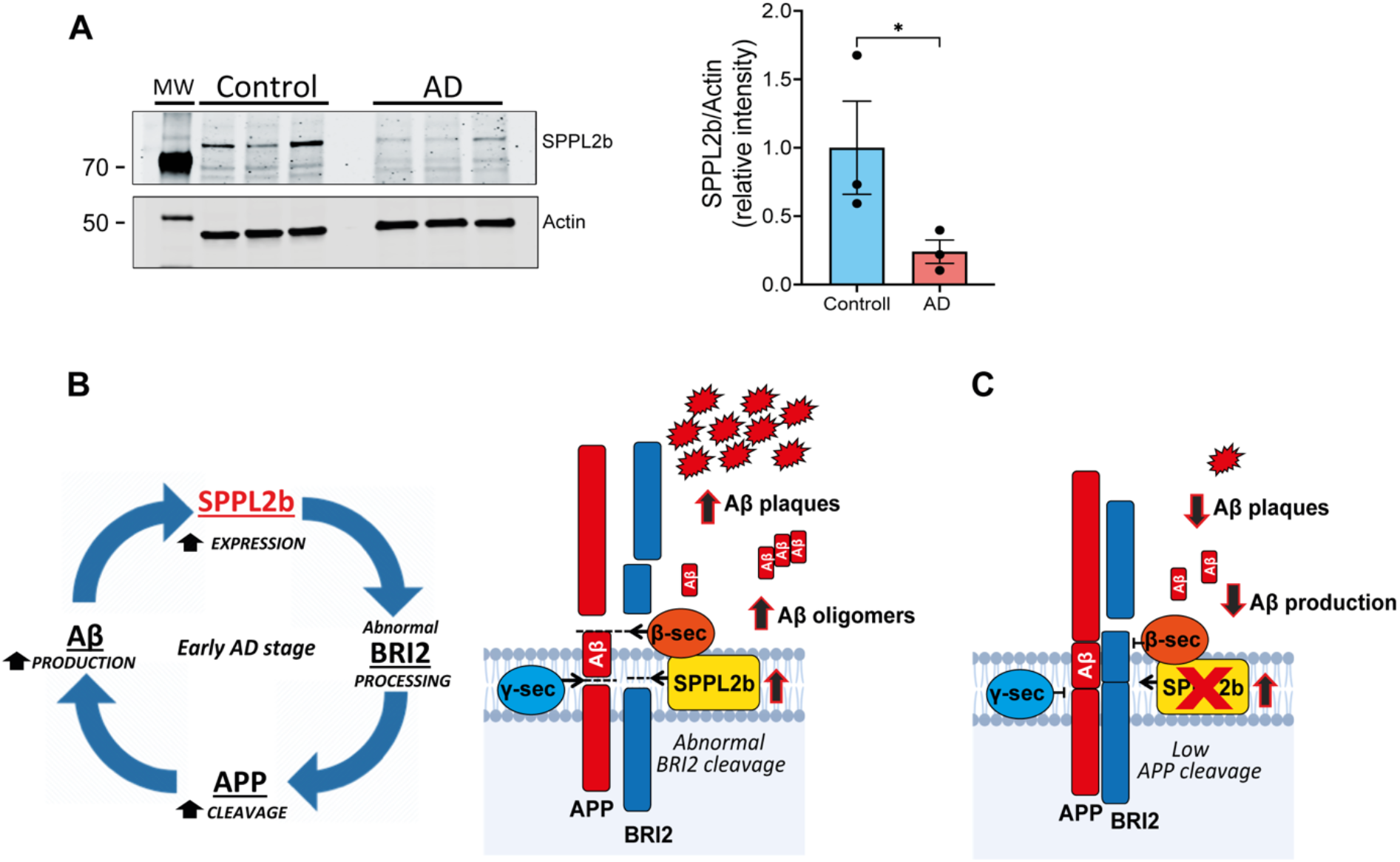
SPPL2b is downregulated in AD human prefrontal cortex. **(A)** Western blot analysis of SPPL2b expression in the human prefrontal cortex of healthy control (Control) and AD cases (AD) from post-mortem tissues in the late stage of the disease (Braak stages V-VI) (n = 3). Data are represented as mean ± S.E.M. *P < 0.05 significantly different from the healthy controls. Data were analyzed by unpaired Student’s t-test. (**B**) Summarizing the results obtained support that Aβ42 is directly involved in SPPL2b expression, potentially generating a vicious cycle where Aβ42, SPPL2b, BRI2, and APP are involved. To the right a schematic model depicting function of SPPL2b in this amyloidogenic pathway, where a high expression of SPPL2b in the early stage of AD affects BRI2 processing resulting in a lower APP-BRI2 interaction and an altered APP cleavage and subsequent increased Aβ production. (**C**) On the other hand, inhibition of SPPL2b activity can restore the physiological condition and significantly decrease the APP cleavage and lower Aβ production.

Taking all together, Aβ42 modulates SPPL2b expression, which correlates with an altered BRI2 detection and APP processing potentially inducing a vicious cycle where Aβ42, SPPL2b, BRI2, and APP are involved (Fig. 8B). Furthermore, we speculate that SPPL2b can play an important role in the onset of AD pathology, where an up-regulation in the early stages can play a crucial function in the AD amyloidogenic pathways (Fig. 8C).

## DISCUSSION

Intramembrane proteolysis is an essential cellular mechanism in several signaling pathways and protein degradation (53). Among intramembrane proteases, the γ-secretase catalytic subunits PS1 and PS2 are involved in AD (54,55). PS1 and PS2, with their analogs SPP and the SPPLs, are part of the GXGD-type aspartyl proteases family (56,57). γ-secretase is directly involved in the cleavage of the APP transmembrane region, and its roles in familiar AD have been well characterized since mutations in PS1 and PS2 are directly implicated in early-onset familial AD. On the other hand, limited information is available regarding the role of SPPLs in AD. Since the discovery of the amyloidogenic pathway, attenuating Aβ production by inhibiting the proteases β- or γ-secretase has been considered an attractive strategy for preventing the disease progression in patients suffering from AD. However, these protease inhibition approaches have met several challenges and limitations over recent years (14). Hence, a better understanding and characterization of new proteins involved in the Aβ cascade may provide precious information and potentially novel AD therapeutic targets.

Towards that end, we here disclose the involvement of SPPL2b in AD pathology using both in vitro and in vivo approaches. SPPL2b has been linked to AD in a previous study in which a higher level of this enzyme was reported in postmortem brain tissues from patients in an early stage of AD (58). In addition, several pieces of evidence connected Bri2, an SPPL2b substrate to AD (59,60). However, the characterization of SPPL2b at the endogenous level in state-of-the-art AD models and neuronal cells was necessary. Our findings demonstrate that the SPPL2b protein levels follow a differential expression pattern during AD pathology, based on the degree of the amyloid pathology which in turn affects the processing of APP. In particular, our data show that SPPL2b expression is increased in the early stage of AD. Conversely, advanced AD stages are associated with a significant SPPL2b downregulation, a circumstance consistently observed in Aβ42-treated cells, *App*^*NL-G-F*^ AD mice, and human AD brains.

SPPL2b is highly expressed in the brain and, mainly, in the hippocampus and cortex (21). One of the substrates of SPPL2b is the transmembrane protein BRI2, for which SPPL2b is responsible for cleaving it in its intramembrane region. Several studies have previously reported that BRI2 interacts with APP and negatively modulates APP cleavage by masking the secretase accesses (38,39,59,61). The BRI2 region involved in the interaction with APP consists of the 46-106 amino acids sequence that includes the transmembrane sequence and the first portion of the extracellular domain (62). Del Campo and colleagues proposed a very intriguing interpretation of how the different processing of BRI2 leads to pro-aggregating or antiaggregating effects, respectively, by preventing or allowing its interaction with APP. Abnormal levels of furin, ADAM10 and SPPL2b in the early AD stage affect the BRI2 cleavage inducing the misfolding of the BRI2 ectodomain, which forms aggregates that facilitate Aβ accumulation and deposition (42,60). Our results support this interpretation; up-regulation of SPPL2b correlates with increased secretion of the BRICHOS-containing fragment. Most importantly, SPPL2b overexpression was associated with a reduction of the mature form of APP while increasing its soluble fraction.

Hence, to determine whether SPPL2b expression is directly affected by Aβ pathology, we have evaluated *in vitro* and *ex vivo* the effect of AD-causing Aβ42. Western blot analysis showed that low doses of Aβ42 induce an up-regulation of SPPL2b in WT cells, a phenomenon also observed in WT ex vivo slices in the cortex. Interestingly, a higher dose of Aβ42 induced a down-regulation of SPPL2b. Consequently, we sought to verify whether the Aβ pathology affects the SPPL2b expression in the AD *App*^*NL-G-F*^ knock-in mouse model. To this end, we measured SPPL2b in the cortex and hippocampus of *App*^*NL-G-F*^ mice during the early Aβ pathology stage (3 months of age) and in an advanced AD stage (10 and 22 months of age), allowing us to verify whether the progression of Aβ pathology and, with, increasing concentration of Aβ42 influenced the levels of SPPL2b in a state-of-the-art AD model (63). Strikingly, the results obtained *in vitro* were confirmed in *App*^*NL-G-F*^ mice. SPPL2b levels were increased in the cortex of 3 months old mice, similar to what we observed in *ex vivo* slices incubated with Aβ42. Wheras, in old mice, which show a marked deposition of Aβ42 aggregates (44,63), SPPL2b was downregulated in both the cortex and hippocampus. Taken together a biphasic effect is observed in the *App*^*NL-G-F*^ mice, with an up-regulation during the early Aβ pathology stage and a down-regulation in an advanced AD stage. The higher levels of SPPL2b during the early stages of AD might lead to altered BRI2 processing and, thus, to a reduced BRI2-APP interaction that allows the secretase to access their cleavage site in APP and increase the production of Aβ and plaque deposition. Those data are in line with the results reported by Del Campo and colleagues (42), who reported a significant increase of SPPL2b in AD postmortem patients at Braak Stage II and III, and this high expression of SPPL2b correlated with an abnormal BRI2 processing and a reduced presence of the APP-BRI2 complexes. This process could likely play an important role in the onset of the disease.

Previous findings proposed that SPPL2b only efficiently cleaves BRI2 after an initial shedding of mBRI2 by ADAM-10 (24,46). However, these data came from cell models in non-AD pathological conditions. We speculate that a different BRI2 shedding (ADAM10-independent) can occur upon high expression of SPPL2b. In addition, taking into consideration also the previous study reporting that all the proteases involved in BRI2 processing have an abnormal expression in AD human hippocampus (42). Furthermore, a non-canonical ectodomain cleavage has also been recently reported for SPPL2a (64). However, further studies are necessary to confirm this hypothesis, which will also consider the possibility of SPPL2b influencing ADAM10/17 expression.

Conversely, in the late stages of AD, when the disease and the Aβ pathology are severe, we observed a reduction of SPPL2b expression. One of the hypothetical explanations for this down-regulation could be motivated by an overstimulation of the system, that in biology is most of the time associated with a reduction of gene expression (65). We also speculate that the initial over-expression of SPPL2b could play a key role in the production and accumulation of Aβ; instead, the reduction of SPPL2b in AD late phases could play an important role in reducing the BRI2 cleavage and increase the APP-BRI2 interaction in an attempt to inhibit the Aβ production. Given the presence of SPPL2b in neuronal spines, another important point to consider is that the reduction of SPPL2b expression in late AD stages could be linked to the reduction of the neuronal spines, previously observed in *App*^*NL-G-F*^ mice (65) since no neuronal loss has been reported in *App*^*NL-G-F*^ mice (44).

Most importantly, the analysis of the media from SPPL2b KO primary neuronal culture strongly supports that inhibition of SPPL2b leads to a reduction of Aβ40 and Aβ42 production.

The biphasic expression of SPPL2b during AD pathology seems to be mainly linked to the SPPL2b expression levels in the neuronal cells. However, SPPL2b was also detected in primary microglial cell culture where its expression increased after Aβ exposition at the dose of 1μM. Consistent with this, high SPPL2b positive staining was observed in microglia surrounding the plaques in old *App*^*NL-G-F*^ mice.

Together with BRI2, TNFα is one of the main substrates of SPPL2b and is highly expressed in microglia cells. During AD, microglia produce increased levels of cytotoxic and inflammatory mediators, such as TNFα, which can start a positive feedback mechanism that reactivates microglia itself. Soluble TNFα is released after ADAM17 cleavage and the remaining TNFα-NTF is processed by SPPL2b. Accordingly, we speculate that high SPPL2b levels are promoting the cleavage of TNFα in its transmembrane domain, increasing the production of TNFα ICD. TNFα ICD has been associated with the production of IL12, a proinflammatory cytokine present in high concentrations in AD and whose pathway, when inhibited, is associated with a significant improvement in AD pathology (12). Finally, SPPL2b expression was also detected in astrocytes, although in a less amount, its expression seems to be not affected in Aβ pathology.

Although the results outlined in this study confirm the hypothesis of an important and relevant relationship between SPPL2b and AD, this study has a major limitation which is the lack of data regarding a potential therapeutic effect of a pharmacological regulation of SPPL2b.

Unfortunately, to date, no specific inhibitor is available for SPPL2b, underlying the lack of selective inhibiting compounds. However, as an imminent future intention, we plan to delete the SPPL2b gene expression in AD mice context to verify whether AD pathology might be reduced (Fig. 8C). Finally, considering that LPS increased SPPL2b expression in the hippocampus of *ex vivo* slices, a link between this protein and general inflammation can not be excluded and should be an object of other more dedicated research.

In conclusion, the results outlined in this study support a relevant connection between SPPL2b and AD. Our findings show that SPPL2b is increased in the early stages of AD, suggesting a potential involvement in the pathogenesis of the disease. In a global scenario characterized by the desperate need to identify novel strategies to prevent and counteract AD progression, this study points out and strengthens the importance of SPPL2b in the Aβ cascade, allowing verifying whether targeting this protein might shed new light on therapeutic approaches for AD.

## SUPPLEMENTARY MATERIAL

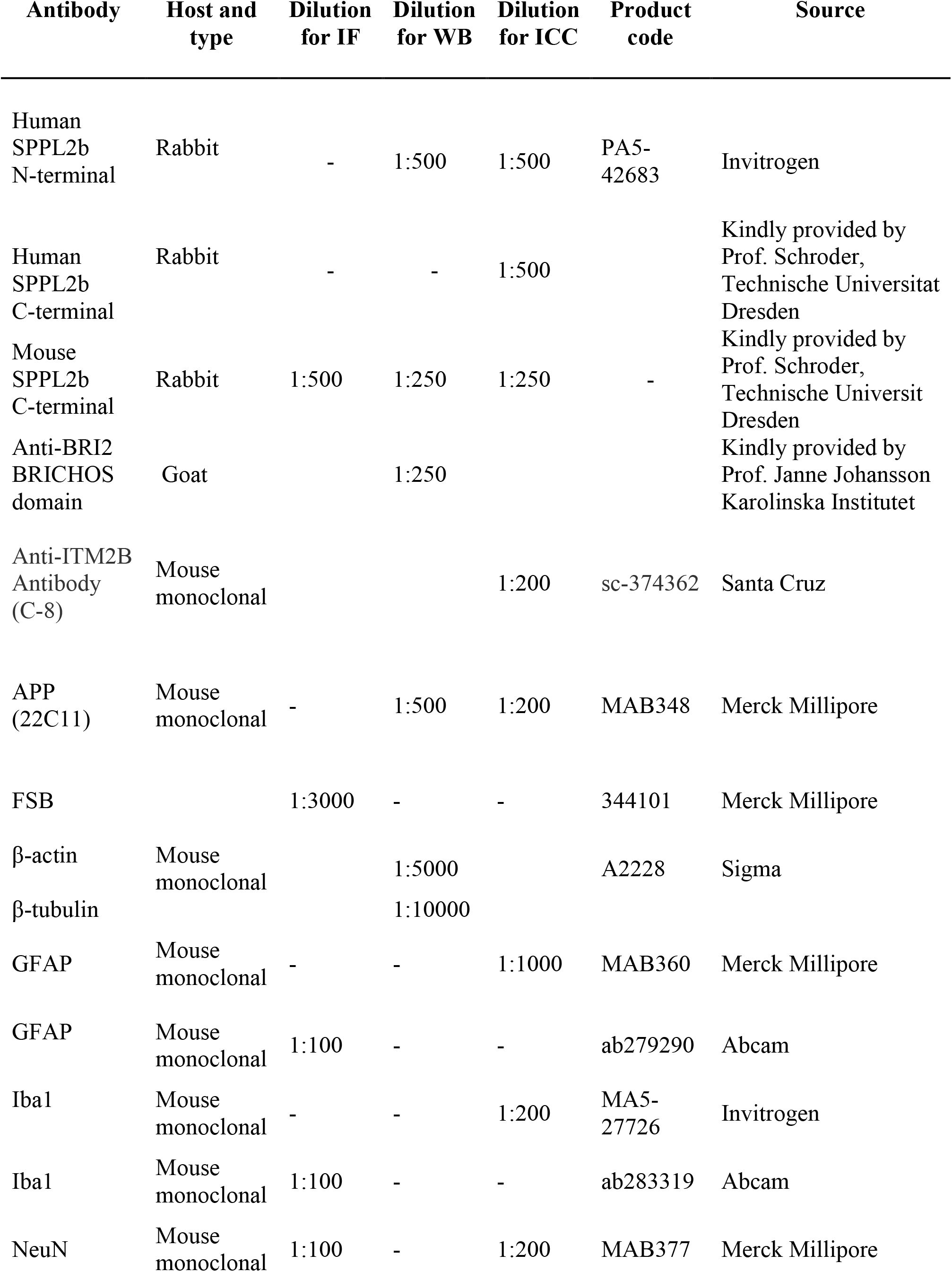
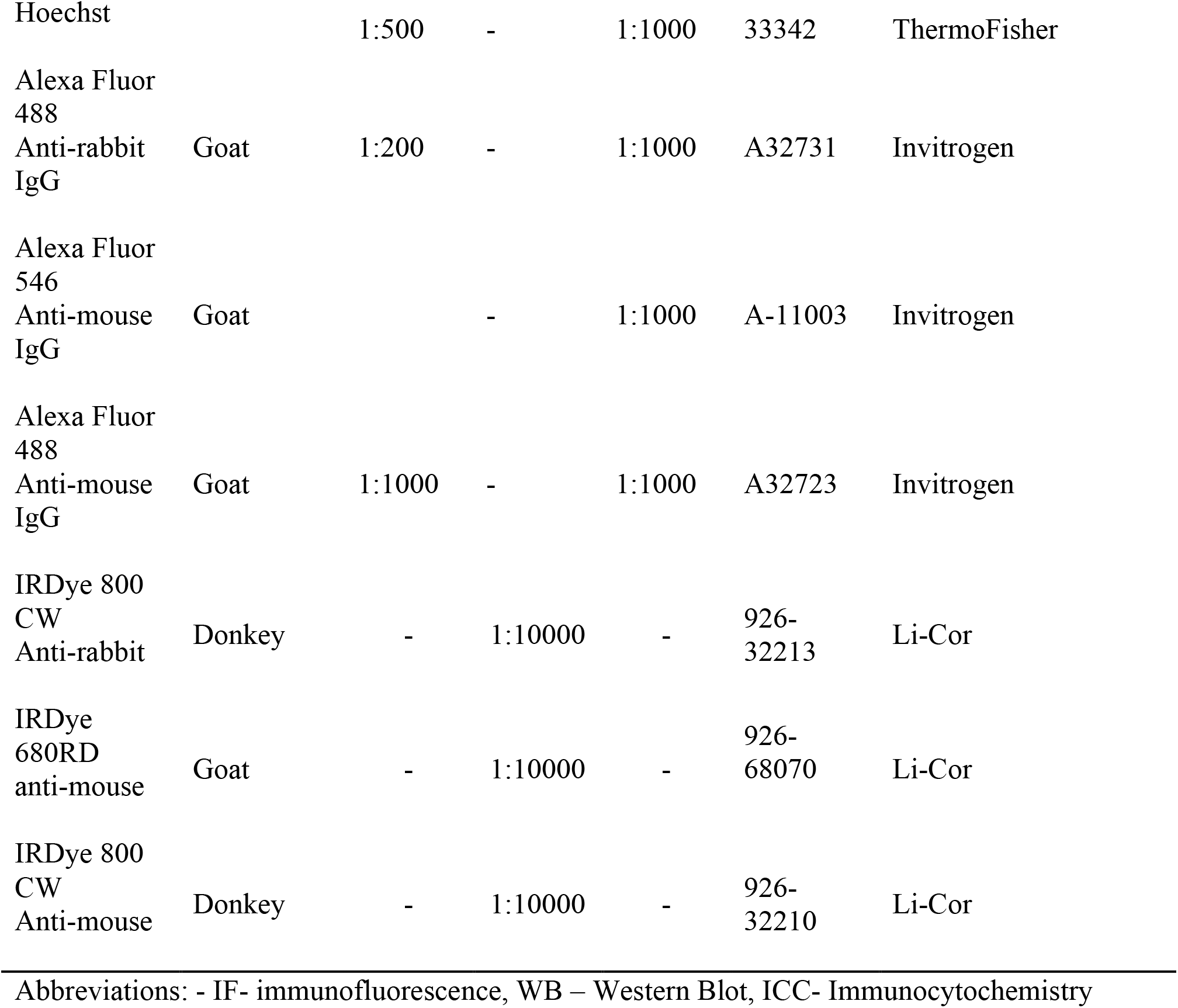

**Figure S1.**
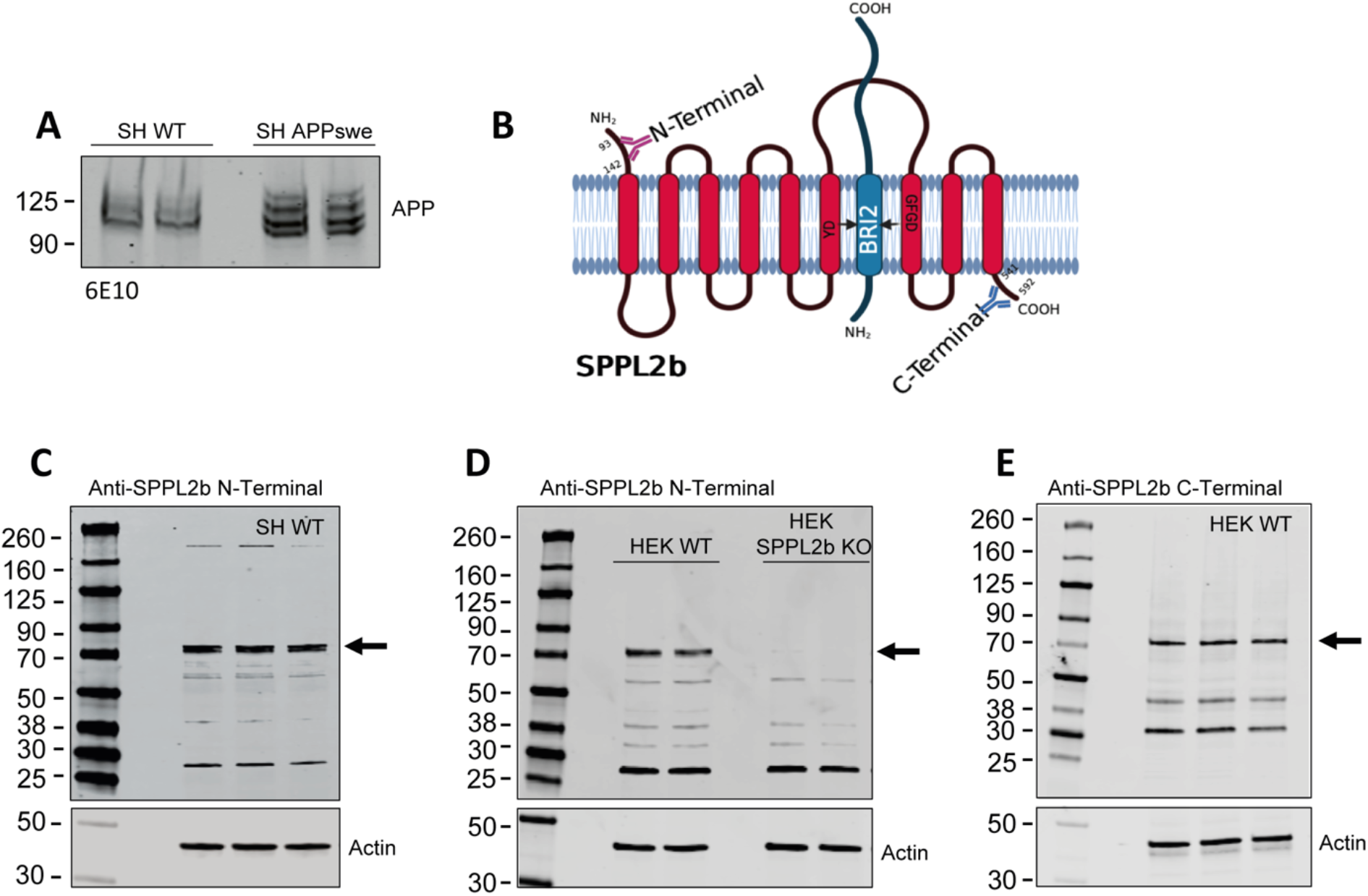
Evaluation of the overexpression of human APPswe and SPPL2b antibodies specificity in the human cell models SH-SY5Y and HEK293. (**A**) Western blot staining of APP (6E10) in SH-SY5Y WT and SH-SY5Y overexpressing APPswe. (**B**) Schematic two-dimensional representation of SPPL2b protein, consisting of nine transmembrane domains, and its catalytic site in presence of SPPL2b substrate BRI2. The epitopes of the anti-SPPL2b N-terminal (93-142 aa) and anti-SPPL2b C-terminal (541-592 aa) antibodies are indicated in the figures (the figure has been prepared by using biorender.com) (**C**) Representative Western blots obtained by using the anti-SPPL2b N-terminal in SH-SY5Y WT cell lysate, and (**D**) in HEK293 WT and HEK293 after SPPL2b gene knock-out. (**E**) Representative Western blot obtained by using the anti-SPPL2b C-terminal in HEK293 WT cell lysate. The arrows in C, D, and E indicate the SPPL2b bands at 70kDa.

**Figure S2.**
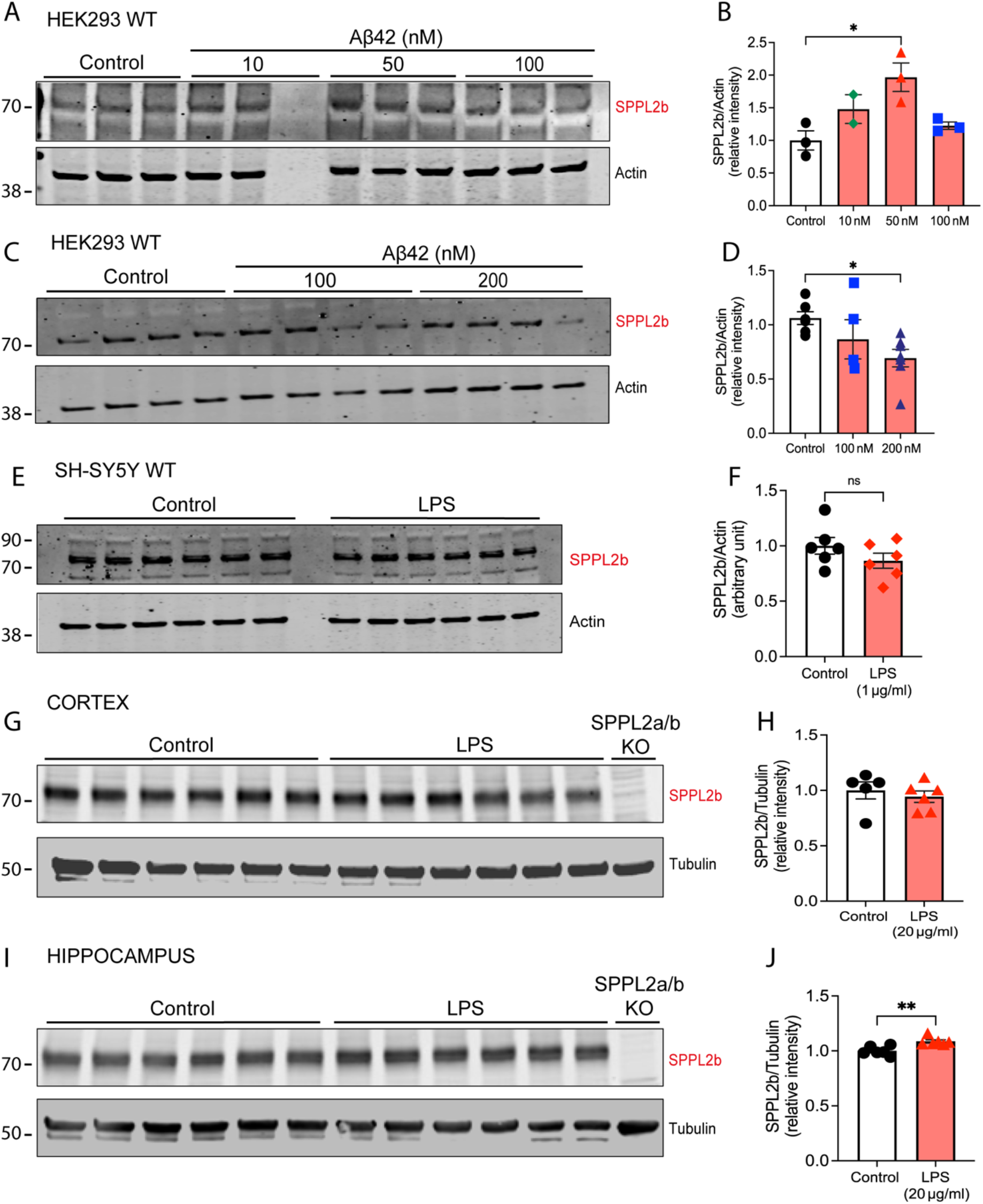
SPPL2b expression in HEK293 WT control cells and after Aβ42 treatment. (**A, B**) Representative Western blot of HEK293 WT treated with Aβ42 (10-100 nM) or (**C, D**) 100 and 200 nM for 24h. (**E, F**) Western blot evaluation of SPPL2b expression in SH-SY5Y WT cells after treatment with LPS (1 μg/ml) for 24 h. (**G, H**) Representative Western blot analysis of mice brain cortex and (**I, J**) hippocampal slices kept ex vivo in artificial CSF and treated with LPS 20 μg/ml for 6 hours. Results are normalized to tubulin protein expression. Data are represented as mean ± S.E.M. **P < 0.01, *P < 0.05 significantly different from the Control. Data were analyzed either by unpaired Student’s t-test (B, H, J) or by one-way ANOVA with Tukey’s multiple comparison test (D, F).

**Figure S3.**
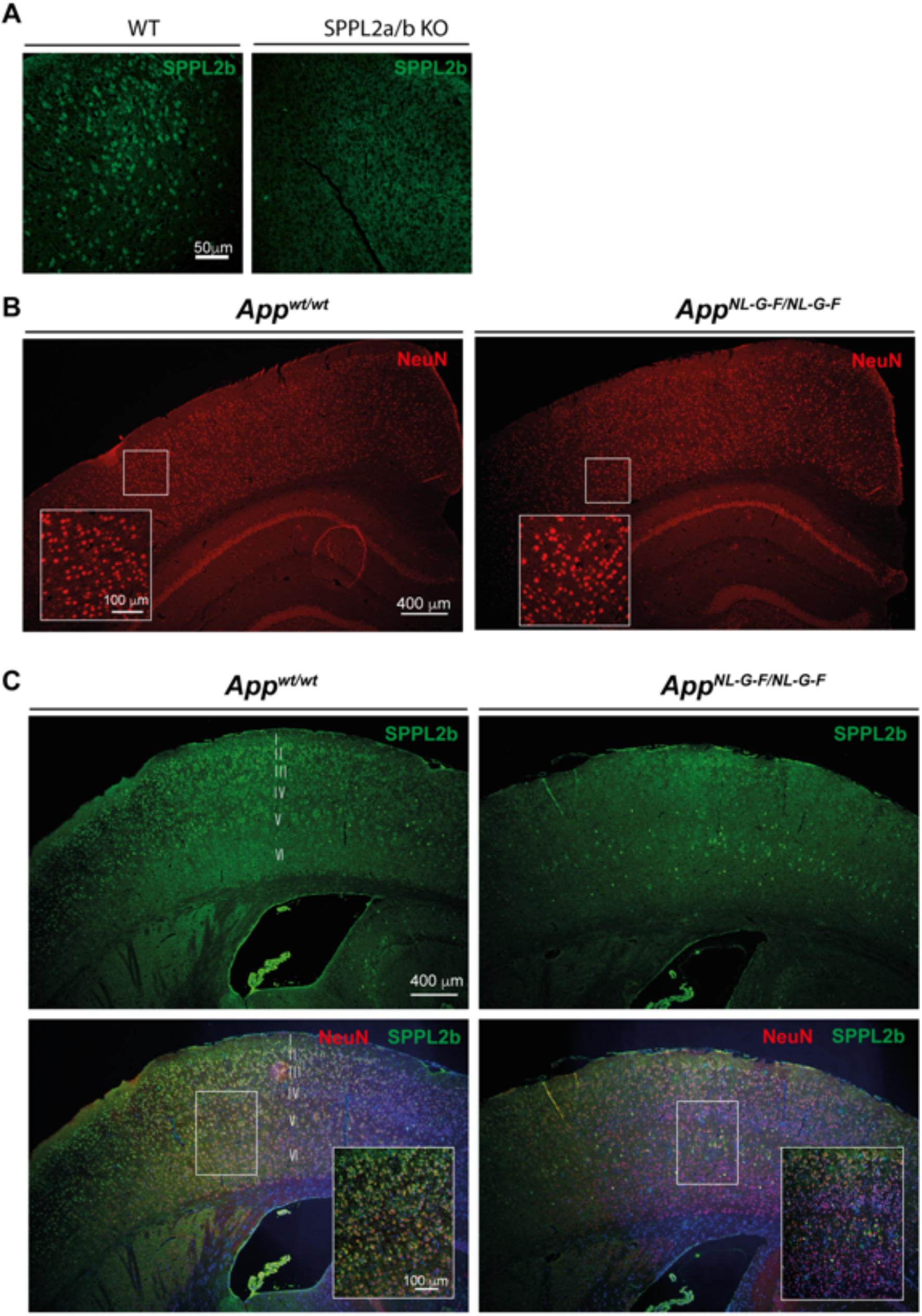
SPPL2b staining in the cortex area of WT and *App*^*NL-G-F*^ mice. (**A**) Representative immunohistochemistry showing SPPL2b staining in WT and SPPL2a/b KO mice. (**B**) Neuronal staining (NeuN) in 10 months old WT and *App*^*NL-G-F*^ mice. (**C**) Representative immunohistochemistry showing SPPL2b staining in 10 months old WT and *App*^*NL-G-F*^ mice. I, II, III, IV, V, and VI indicate the cortex layers. Scale bar sizes are reported in the figure.

**Figure S4.**
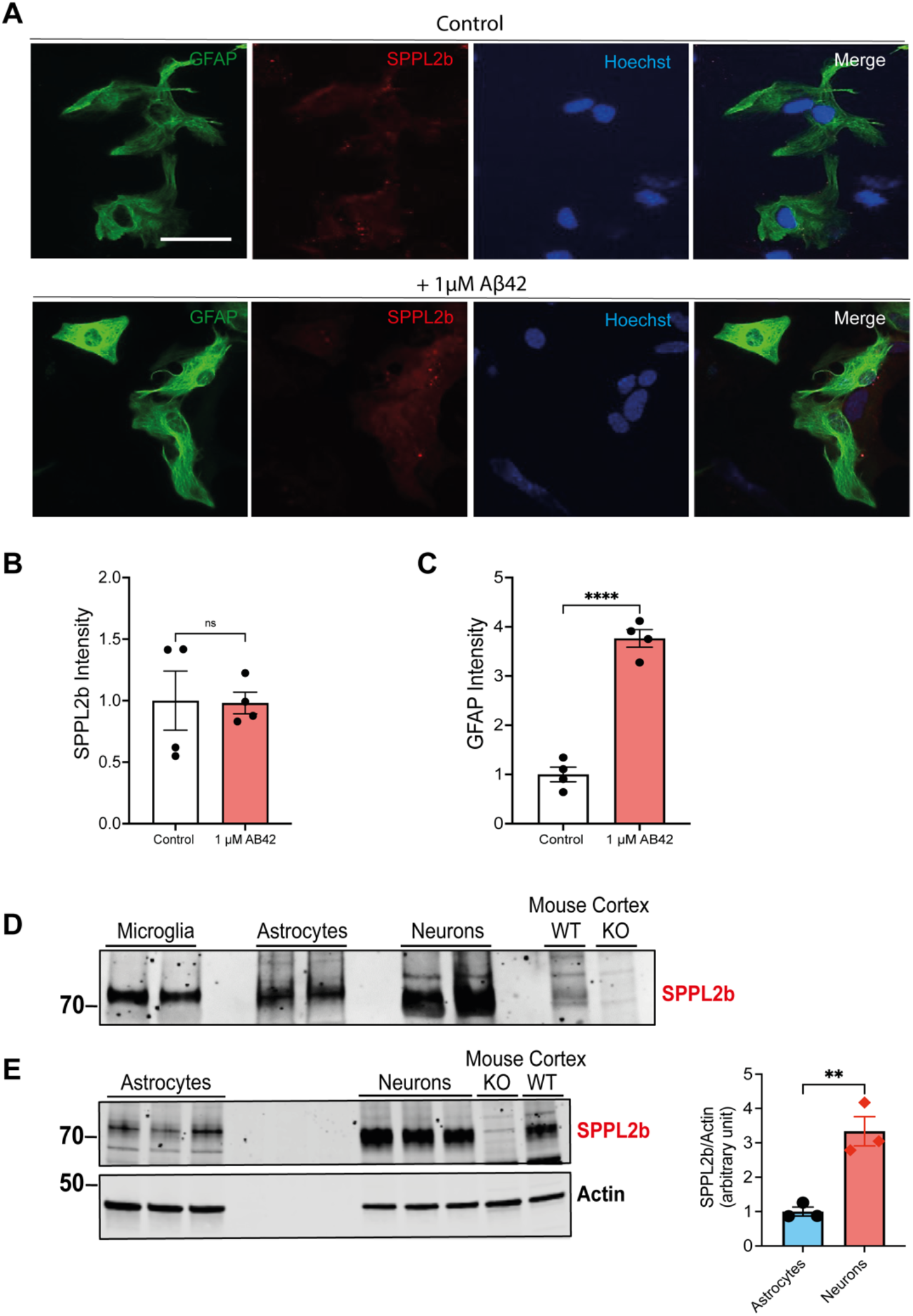
SPPL2b expression levels in primary astrocyte cells before and after Aβ42 treatment. (**A**) Astroglia (GFAP, green), SPPL2b (red), and nucleus (Hoechst blue) stainings in astrocyte cells non-treated (control) and after 24-hour treatment with 1 μM Aβ42. Scale bar: 20 μm. (**B, C**) Quantification of GFAP and SPPL2b staining. (**D**) Representative Western blot of SPPL2b in microglia, astrocytes, and neuronal cells from isolated primary cell culture of WT mice. (E) Representative Western blot showing the SPPL2b expression in astrocytes and neurons and quantification (n=3). In D and E, the last 2 lanes have been loaded WT and SPPL2a/b KO mouse cortex samples as a positive and negative control, respectively. Data are represented as mean ± S.E.M. ****P < 0.0001, **P < 0.01, significantly different from the control. Data were analyzed either by unpaired Student’s t-test

## Acknowledgments

The authors acknowledge Takaomi Saido and Takashi Saito at the RIKEN Center for Brain Science for providing App knockin mice, Dr. Janne Johansson (Karolinska Instituet, Sweden) for antibody donation.

This work was supported by the Olle Engkvists Stiftelse, Gun & Bertil Stohnes Stiftelse, Demensfonden, Lindhés Advokatbyrå Stiftelse, Åhlén-stiftelsen, Gamla tjänarinnor, Alzheimerfonden, Hållsten Research Foundation, Sonja Leikrans donation, as well as by the Research Group FOR2290 of the Deutsche Forschungsgemeinschaft as well as by the grants FL 635/2-2 and FL 635/2-3.

## Author contributions

RM., CT, and SZ. performed experiments, analyzed data, and contribute to writing the paper. AK, YAT, FP, CG, GC, LL, AF, and RJ, performed experiments and analyzed data. RF, TM, and BS supervised experiments and critically revised the paper. PN planned, and supervised experiments and contributes to writing the paper. ST conceptualized the study, planned, and supervised experiments, analyzed data, and wrote the paper. All authors commented on the manuscript.

## Data and materials availability

All data needed to evaluate the conclusions in the paper are present in the paper and/or the Supplementary Materials.

